# Histone proteoform analysis reveals epigenetic changes in adult mouse brown adipose tissue in response to cold stress

**DOI:** 10.1101/2023.07.30.551059

**Authors:** Bethany C. Taylor, Loic H. Steinthal, Michelle Dias, Hari K. Yalamanchili, Scott A. Ochsner, Gladys E. Zapata, Nitesh R. Mehta, Neil J. McKenna, Nicolas L. Young, Alli M. Nuotio-Antar

## Abstract

Regulation of the thermogenic response by brown adipose tissue (BAT) is an important component of energy homeostasis with implications for the treatment of obesity and diabetes. Our preliminary analyses uncovered many nodes representing epigenetic modifiers that are altered in BAT in response to chronic thermogenic activation. Thus, we hypothesized that chronic thermogenic activation broadly alters epigenetic modifications of DNA and histones in BAT. Motivated to understand how BAT function is regulated epigenetically, we developed a novel method for the first-ever unbiased top-down proteomic quantitation of histone modifications in BAT and validated our results with a multi-omic approach. To test our hypothesis, wildtype male C57BL/6J mice were housed under chronic conditions of thermoneutral temperature (TN, 28.8°C), mild cold/room temperature (RT, 22°C), or severe cold (SC, 8°C) and BAT was analyzed for DNA methylation and histone modifications. Methylation of promoters and intragenic regions in genomic DNA decrease in response to chronic cold exposure. Integration of DNA methylation and RNA expression data suggest a role for epigenetic modification of DNA in gene regulation in response to cold. In response to cold housing, we observe increased bulk acetylation of histones H3.2 and H4, increased histone H3.2 proteoforms with di- and trimethylation of lysine 9 (K9me2 and K9me3), and increased histone H4 proteoforms with acetylation of lysine 16 (K16ac) in BAT. Taken together, our results reveal global epigenetically-regulated transcriptional “on” and “off” signals in murine BAT in response to varying degrees of chronic cold stimuli and establish a novel methodology to quantitatively study histones in BAT, allowing for direct comparisons to decipher mechanistic changes during the thermogenic response. Additionally, we make histone PTM and proteoform quantitation, RNA splicing, RRBS, and transcriptional footprint datasets available as a resource for future research.

## Introduction

Increasing activation of brown adipose tissue (BAT) has been suggested as a therapy to counter obesity and type 2 diabetes (Cypess et al., 2009; Cypess and Kahn, 2010). BAT is a mitochondria-rich tissue that is essential for maintaining body temperature through nonshivering thermogenesis (NST). During thermogenic activation, uptake and oxidation of circulating glucose and fatty acid into BAT renders the tissue a biologic “sink” for both nutrients (Labbé et al., 2015). Altered BAT function in adults is associated with differences in adiposity, glucose homeostasis, insulin sensitivity, and energy expenditure (Chondronikola et al., 2014; Cypess et al., 2009; Saito et al., 2009; van Marken Lichtenbelt et al., 2009). In particular, NST-induced glucose uptake and oxidation by BAT is attenuated in obese and type 2 diabetic patients (Blondin et al., 2015). While prior studies have established that specific enzymes that modulate DNA methylation and histone modifications are required for the proper differentiation and development of BAT, quantitative data and rigorous comparisons regarding global DNA methylation changes and histone modifications in BAT in response to varying degrees of thermogenic activation in adults is lacking (Nic-Can et al., 2019; Xiao and Kang, 2019; Yi et al., 2020). Thus, epigenetic regulation of the thermogenic response in adult BAT remains mostly unexplored and may yield insights into means through which tissue function and consequent beneficial effects on whole-body metabolism may be restored.

Histone modifications are particularly less studied in BAT due to a lack of effective methods for the isolation of nuclei from BAT, acid extraction from fatty tissues, and appropriate methods of analysis (Strzyz, 2016). Here, we have developed a method for the first unbiased, top-down mass spectrometry-based proteomic analysis of histone modifications in murine BAT depots. This method allows us to quantitate discrete post-translational modifications (PTMs) and proteoforms of histones H4 and H3 that are associated with different thermogenic responses in BAT from adult mice for the first time. Using this method, reduced representation bisulfite sequencing (RRBS), and findings from RNA-Seq data, we test our hypothesis that the thermogenic response in adult BAT is associated with multiple epigenetic changes that impact gene expression. We integrate our findings for the first ever comprehensive and unbiased analysis of epigenetic changes associated specifically with BAT thermogenic activation.

## Results

### Analysis of RNA-Seq data reveals housing temperature-dependent effects on genes involved in epigenetic regulation

Acclimation and BAT remodeling in response to ambient temperatures occurs when mice are housed under constant temperatures for at least 7 days (Virtue and Vidal-Puig, 2013). Additionally, prior studies have established that standard laboratory housing of mice at 20-24°C is a chronic mild cold condition that stimulates NST (Cannon and Nedergaard, 2004; Sanchez-Gurmaches et al., 2018; Virtue and Vidal-Puig, 2013). We reanalyzed and further interrogated publicly-available RNA-Seq data and noted significant cold housing temperature-induced reductions in the expression of genes encoding DNA methyltransferases, *Dnmt1*, *Dnmt3a*, and *Dnmt3b*, acetyl-CoA-generating enzymes, and histone-modifying enzymes *Smyd3, Kdm2b, Setd7, Kdm3a, Kdm5b, Kdm5c*, and *Kdm8* (**Table 2**) (Sanchez-Gurmaches et al., 2018). Expression of ten-eleven translocation (TET) enzymes, *Tet2* and *Tet3*, which facilitate demethylation of DNA 5-methylcytosine, did not significantly differ between TN vs SC conditions (data not shown) (Pastor et al., 2013). In addition, we noted significantly altered expression of genes involved in one-carbon metabolism, suggesting that there may be housing temperature-dependent effects on the methyl donor pool that could also contribute to epigenetic modifications in BAT.

**Table 1.**
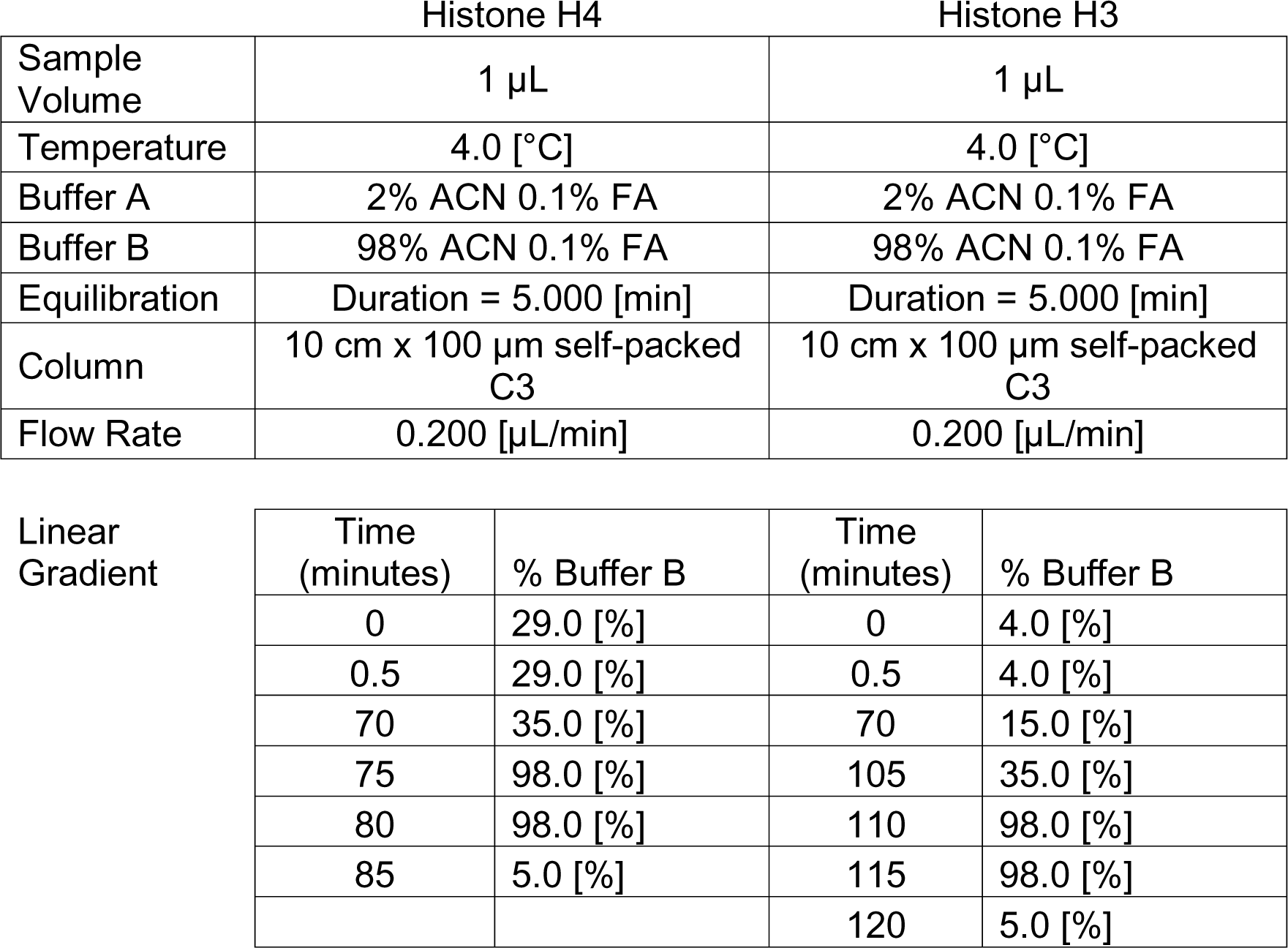
Online HPLC parameters for histones H4 and H3. 1 µL of sample was injected per run after resuspension to a standard concentration (200 ng/µL for H4; 2 µg/µL for H3). The same buffers were used with different gradients (See Linear Gradient section). Histone species were introduced to the Orbitrap Fusion Lumos mass spectrometer using nano-electrospray ionization as they eluted.

|  | Histone H4 | Histone H3 |
| --- | --- | --- |
| Sample Volume | 1 $\mu\text{L}$ | 1 $\mu\text{L}$ |
| Temperature | 4.0 [°C] | 4.0 [°C] |
| Buffer A | 2% ACN 0.1% FA | 2% ACN 0.1% FA |
| Buffer B | 98% ACN 0.1% FA | 98% ACN 0.1% FA |
| Equilibration | Duration = 5.000 [min] | Duration = 5.000 [min] |
| Column | 10 cm x 100 $\mu\text{m}$ self-packed C3 | 10 cm x 100 $\mu\text{m}$ self-packed C3 |
| Flow Rate | 0.200 [$\mu\text{L}/\text{min}$] | 0.200 [$\mu\text{L}/\text{min}$] |

|  |  |  |  |  |
| --- | --- | --- | --- | --- |
| Linear Gradient | Time (minutes) | % Buffer B | Time (minutes) | % Buffer B |
|  | 0 | 29.0 [%] | 0 | 4.0 [%] |
|  | 0.5 | 29.0 [%] | 0.5 | 4.0 [%] |
|  | 70 | 35.0 [%] | 70 | 15.0 [%] |
|  | 75 | 98.0 [%] | 105 | 35.0 [%] |
|  | 80 | 98.0 [%] | 110 | 98.0 [%] |
|  | 85 | 5.0 [%] | 115 | 98.0 [%] |
|  |  |  | 120 | 5.0 [%] |

**Table 2.**
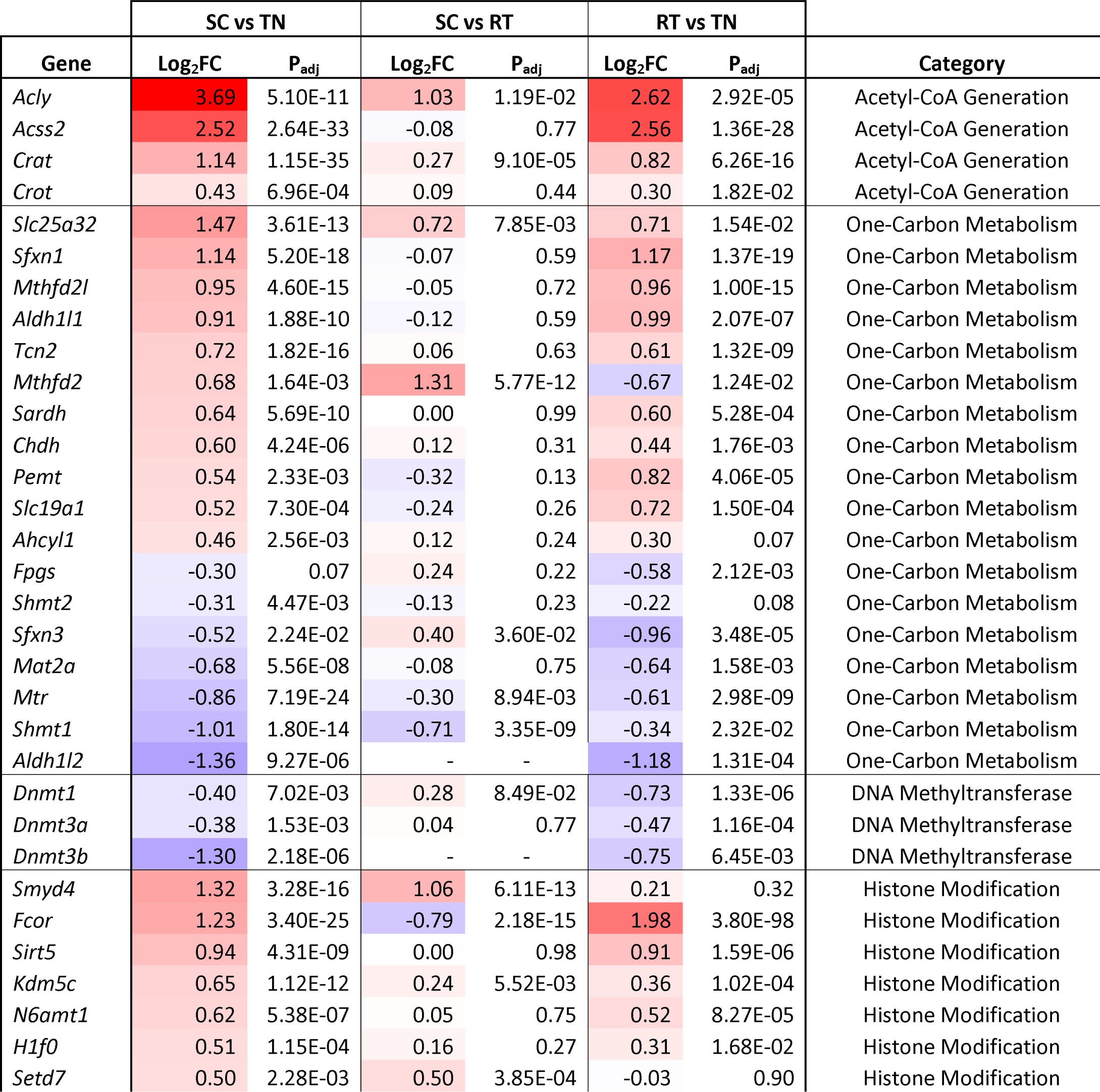
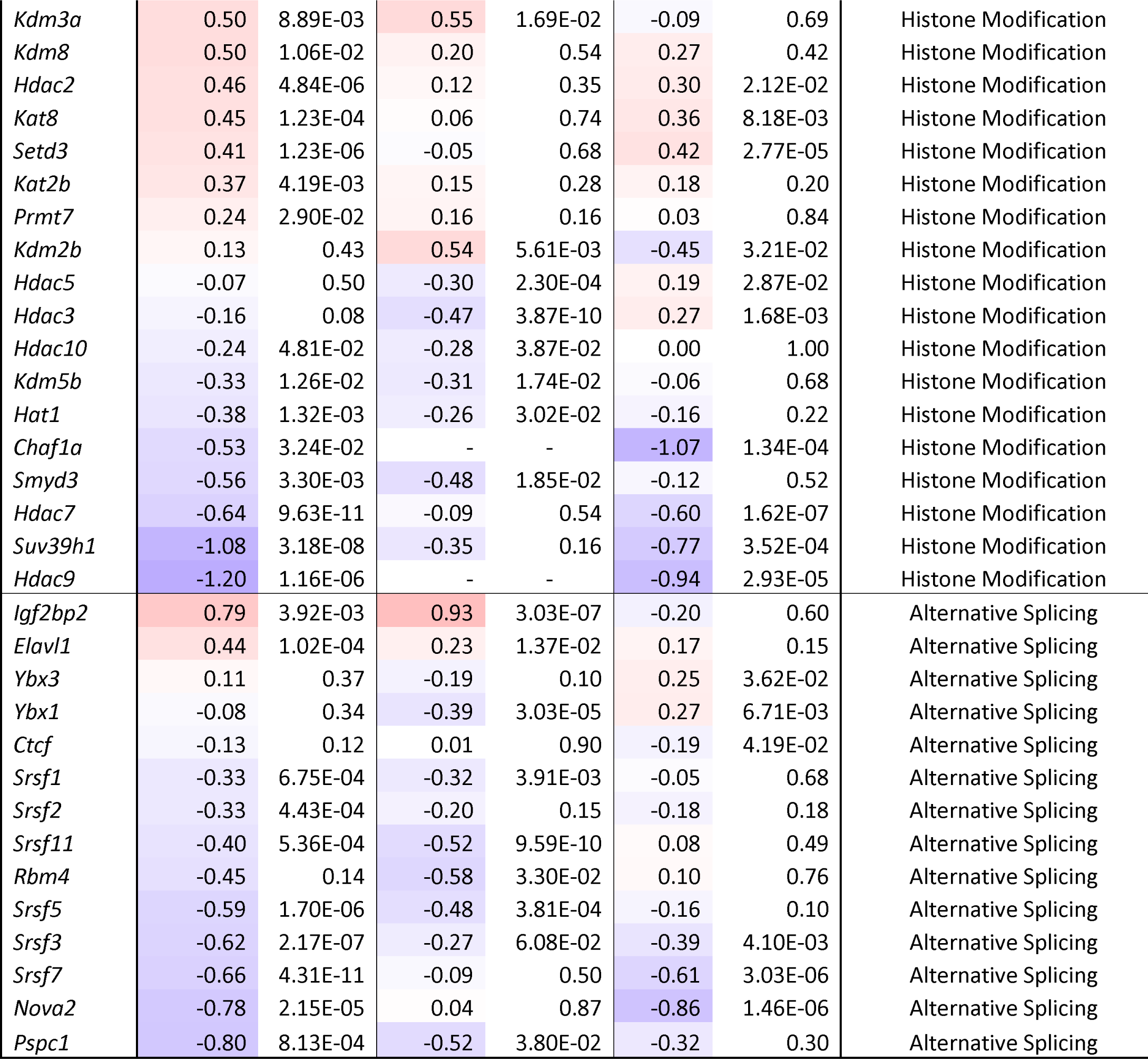
Expression of acetyl-CoA generating enzymes, one-carbon metabolic genes, epigenetic modifiers, and alternative splicing factors is altered in response to chronic thermogenic activation as measured by RNA-seq. TN: thermoneutral; RT: room temperature; SC: severe cold.

|  | SC vs TN |  | SC vs RT |  | RT vs TN |  |  |
| --- | --- | --- | --- | --- | --- | --- | --- |
| Gene | Log<sub>2</sub>FC | P<sub>adj</sub> | Log<sub>2</sub>FC | P<sub>adj</sub> | Log<sub>2</sub>FC | P<sub>adj</sub> | Category |
| *Acly* | 3.69 | 5.10E-11 | 1.03 | 1.19E-02 | 2.62 | 2.92E-05 | Acetyl-CoA Generation |
| *Acss2* | 2.52 | 2.64E-33 | -0.08 | 0.77 | 2.56 | 1.36E-28 | Acetyl-CoA Generation |
| *Crat* | 1.14 | 1.15E-35 | 0.27 | 9.10E-05 | 0.82 | 6.26E-16 | Acetyl-CoA Generation |
| *Crot* | 0.43 | 6.96E-04 | 0.09 | 0.44 | 0.30 | 1.82E-02 | Acetyl-CoA Generation |
| *Slc25a32* | 1.47 | 3.61E-13 | 0.72 | 7.85E-03 | 0.71 | 1.54E-02 | One-Carbon Metabolism |
| *Sfxn1* | 1.14 | 5.20E-18 | -0.07 | 0.59 | 1.17 | 1.37E-19 | One-Carbon Metabolism |
| *Mthfd2l* | 0.95 | 4.60E-15 | -0.05 | 0.72 | 0.96 | 1.00E-15 | One-Carbon Metabolism |
| *Aldh1l1* | 0.91 | 1.88E-10 | -0.12 | 0.59 | 0.99 | 2.07E-07 | One-Carbon Metabolism |
| *Tcn2* | 0.72 | 1.82E-16 | 0.06 | 0.63 | 0.61 | 1.32E-09 | One-Carbon Metabolism |
| *Mthfd2* | 0.68 | 1.64E-03 | 1.31 | 5.77E-12 | -0.67 | 1.24E-02 | One-Carbon Metabolism |
| *Sardh* | 0.64 | 5.69E-10 | 0.00 | 0.99 | 0.60 | 5.28E-04 | One-Carbon Metabolism |
| *Chdh* | 0.60 | 4.24E-06 | 0.12 | 0.31 | 0.44 | 1.76E-03 | One-Carbon Metabolism |
| *Pemt* | 0.54 | 2.33E-03 | -0.32 | 0.13 | 0.82 | 4.06E-05 | One-Carbon Metabolism |
| *Slc19a1* | 0.52 | 7.30E-04 | -0.24 | 0.26 | 0.72 | 1.50E-04 | One-Carbon Metabolism |
| *Ahcyl1* | 0.46 | 2.56E-03 | 0.12 | 0.24 | 0.30 | 0.07 | One-Carbon Metabolism |
| *Fpgs* | -0.30 | 0.07 | 0.24 | 0.22 | -0.58 | 2.12E-03 | One-Carbon Metabolism |
| *Shmt2* | -0.31 | 4.47E-03 | -0.13 | 0.23 | -0.22 | 0.08 | One-Carbon Metabolism |
| *Sfxn3* | -0.52 | 2.24E-02 | 0.40 | 3.60E-02 | -0.96 | 3.48E-05 | One-Carbon Metabolism |
| *Mat2a* | -0.68 | 5.56E-08 | -0.08 | 0.75 | -0.64 | 1.58E-03 | One-Carbon Metabolism |
| *Mtr* | -0.86 | 7.19E-24 | -0.30 | 8.94E-03 | -0.61 | 2.98E-09 | One-Carbon Metabolism |
| *Shmt1* | -1.01 | 1.80E-14 | -0.71 | 3.35E-09 | -0.34 | 2.32E-02 | One-Carbon Metabolism |
| *Aldh1l2* | -1.36 | 9.27E-06 | - | - | -1.18 | 1.31E-04 | One-Carbon Metabolism |
| *Dnmt1* | -0.40 | 7.02E-03 | 0.28 | 8.49E-02 | -0.73 | 1.33E-06 | DNA Methyltransferase |
| *Dnmt3a* | -0.38 | 1.53E-03 | 0.04 | 0.77 | -0.47 | 1.16E-04 | DNA Methyltransferase |
| *Dnmt3b* | -1.30 | 2.18E-06 | - | - | -0.75 | 6.45E-03 | DNA Methyltransferase |
| *Smyd4* | 1.32 | 3.28E-16 | 1.06 | 6.11E-13 | 0.21 | 0.32 | Histone Modification |
| *Fcor* | 1.23 | 3.40E-25 | -0.79 | 2.18E-15 | 1.98 | 3.80E-98 | Histone Modification |
| *Sirt5* | 0.94 | 4.31E-09 | 0.00 | 0.98 | 0.91 | 1.59E-06 | Histone Modification |
| *Kdm5c* | 0.65 | 1.12E-12 | 0.24 | 5.52E-03 | 0.36 | 1.02E-04 | Histone Modification |
| *N6amt1* | 0.62 | 5.38E-07 | 0.05 | 0.75 | 0.52 | 8.27E-05 | Histone Modification |
| *H1f0* | 0.51 | 1.15E-04 | 0.16 | 0.27 | 0.31 | 1.68E-02 | Histone Modification |
| *Setd7* | 0.50 | 2.28E-03 | 0.50 | 3.85E-04 | -0.03 | 0.90 | Histone Modification |
| *Kdm3a* | 0.50 | 8.89E-03 | 0.55 | 1.69E-02 | -0.09 | 0.69 | Histone Modification |
| *Kdm8* | 0.50 | 1.06E-02 | 0.20 | 0.54 | 0.27 | 0.42 | Histone Modification |
| *Hdac2* | 0.46 | 4.84E-06 | 0.12 | 0.35 | 0.30 | 2.12E-02 | Histone Modification |
| *Kat8* | 0.45 | 1.23E-04 | 0.06 | 0.74 | 0.36 | 8.18E-03 | Histone Modification |
| *Setd3* | 0.41 | 1.23E-06 | -0.05 | 0.68 | 0.42 | 2.77E-05 | Histone Modification |
| *Kat2b* | 0.37 | 4.19E-03 | 0.15 | 0.28 | 0.18 | 0.20 | Histone Modification |
| *Prmt7* | 0.24 | 2.90E-02 | 0.16 | 0.16 | 0.03 | 0.84 | Histone Modification |
| *Kdm2b* | 0.13 | 0.43 | 0.54 | 5.61E-03 | -0.45 | 3.21E-02 | Histone Modification |
| *Hdac5* | -0.07 | 0.50 | -0.30 | 2.30E-04 | 0.19 | 2.87E-02 | Histone Modification |
| *Hdac3* | -0.16 | 0.08 | -0.47 | 3.87E-10 | 0.27 | 1.68E-03 | Histone Modification |
| *Hdac10* | -0.24 | 4.81E-02 | -0.28 | 3.87E-02 | 0.00 | 1.00 | Histone Modification |
| *Kdm5b* | -0.33 | 1.26E-02 | -0.31 | 1.74E-02 | -0.06 | 0.68 | Histone Modification |
| *Hat1* | -0.38 | 1.32E-03 | -0.26 | 3.02E-02 | -0.16 | 0.22 | Histone Modification |
| *Chaf1a* | -0.53 | 3.24E-02 | - | - | -1.07 | 1.34E-04 | Histone Modification |
| *Smyd3* | -0.56 | 3.30E-03 | -0.48 | 1.85E-02 | -0.12 | 0.52 | Histone Modification |
| *Hdac7* | -0.64 | 9.63E-11 | -0.09 | 0.54 | -0.60 | 1.62E-07 | Histone Modification |
| *Suv39h1* | -1.08 | 3.18E-08 | -0.35 | 0.16 | -0.77 | 3.52E-04 | Histone Modification |
| *Hdac9* | -1.20 | 1.16E-06 | - | - | -0.94 | 2.93E-05 | Histone Modification |
| *Igf2bp2* | 0.79 | 3.92E-03 | 0.93 | 3.03E-07 | -0.20 | 0.60 | Alternative Splicing |
| *Elavl1* | 0.44 | 1.02E-04 | 0.23 | 1.37E-02 | 0.17 | 0.15 | Alternative Splicing |
| *Ybx3* | 0.11 | 0.37 | -0.19 | 0.10 | 0.25 | 3.62E-02 | Alternative Splicing |
| *Ybx1* | -0.08 | 0.34 | -0.39 | 3.03E-05 | 0.27 | 6.71E-03 | Alternative Splicing |
| *Ctcf* | -0.13 | 0.12 | 0.01 | 0.90 | -0.19 | 4.19E-02 | Alternative Splicing |
| *Srsf1* | -0.33 | 6.75E-04 | -0.32 | 3.91E-03 | -0.05 | 0.68 | Alternative Splicing |
| *Srsf2* | -0.33 | 4.43E-04 | -0.20 | 0.15 | -0.18 | 0.18 | Alternative Splicing |
| *Srsf11* | -0.40 | 5.36E-04 | -0.52 | 9.59E-10 | 0.08 | 0.49 | Alternative Splicing |
| *Rbm4* | -0.45 | 0.14 | -0.58 | 3.30E-02 | 0.10 | 0.76 | Alternative Splicing |
| *Srsf5* | -0.59 | 1.70E-06 | -0.48 | 3.81E-04 | -0.16 | 0.10 | Alternative Splicing |
| *Srsf3* | -0.62 | 2.17E-07 | -0.27 | 6.08E-02 | -0.39 | 4.10E-03 | Alternative Splicing |
| *Srsf7* | -0.66 | 4.31E-11 | -0.09 | 0.50 | -0.61 | 3.03E-06 | Alternative Splicing |
| *Nova2* | -0.78 | 2.15E-05 | 0.04 | 0.87 | -0.86 | 1.46E-06 | Alternative Splicing |
| *Pspc1* | -0.80 | 8.13E-04 | -0.52 | 3.80E-02 | -0.32 | 0.30 | Alternative Splicing |

### Regulatory network analysis resolves epigenetic writer footprints in cold challenge-regulated gene sets

High confidence transcriptional target (HCT) intersection analysis computes gene sets of interest against libraries of high confidence direct targets derived from public ChIP-Seq datasets to resolve regulatory footprints within those gene sets for transcription factors and other signaling nodes (Bissig-Choisat et al., 2021; Chen et al., 2022; Ochsner et al., 2023, 2020; Zapata et al., 2021). To identify signaling nodes whose gain or loss of function contributed to the observed cold challenge expression profiles, we subjected genes significantly induced or repressed in each of the three contrasts to HCT intersection analysis (**Supplemental File S1**). As validation, we reasoned that our analysis should identify strong footprints within cold exposure-induced genes for nodes encoded by genes whose deletion in the mouse results in deficient thermoregulatory processes. To test this hypothesis, we retrieved a set of nodes (n=15) mapped to the null phenotype “impaired adaptive thermogenesis” (MP:0011049) or “abnormal circadian temperature homeostasis” (MP:0011020) from the Mouse Genome Database and designated these “thermoregulatory nodes.” We then evaluated the distribution of these thermoregulatory nodes among nodes that had strong regulatory footprints among cold challenge-induced genes in the three contrasts (**Figure 1A-C**) (Blake et al., 2021). Consistent with the reliability of our analysis, we observe strong enrichment of the thermoregulatory nodes among the top ranked cold-induced footprint nodes in all three experiments. These nodes include the nuclear receptors *Pparg*, *Ppara*, and *Esrra*, in addition to *Cebpb* and *Prdm16*, among others (Ahmadian et al., 2011; Carmona et al., 2005; Harms et al., 2014; Shen et al., 2020; Villena et al., 2007).

**Figure 1.**
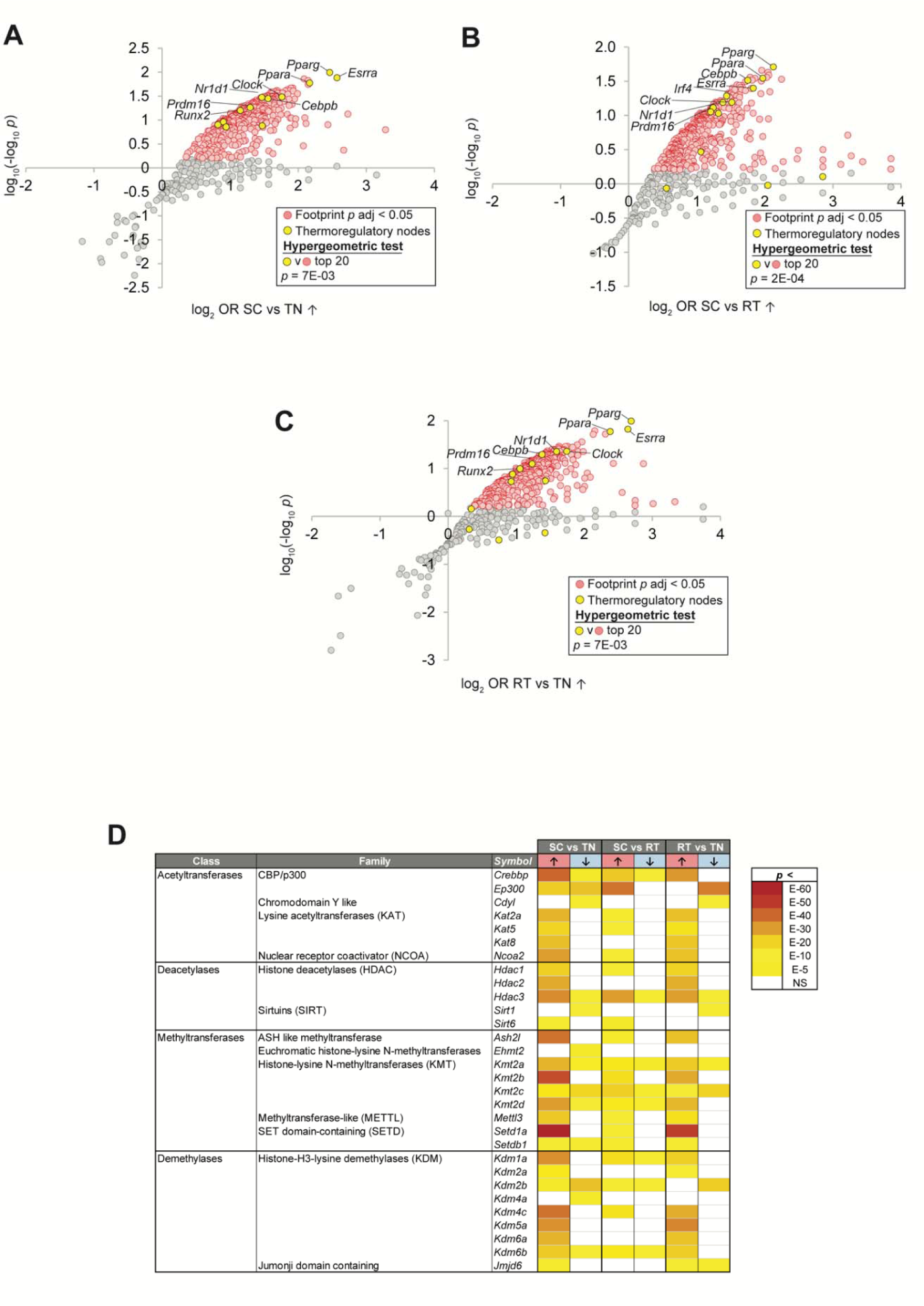
High confidence transcriptional target (HCT) intersection analysis resolves epigenetic writer transcriptional footprints in cold challenge-regulated gene sets. In panels a-c, nodes that have the strongest (higher odds ratio, OR) and most significant (lower p-value) footprints within the indicated cold challenge-induced gene set are distributed towards the upper right of the plot. Scatterplot showing enrichment of nodes with established roles in thermal regulation among nodes that have the most significant intersections with **(a)** SC vs TN-induced genes; **(b)** SC vs RT-induced genes; and **(c)** RT vs TN-induced genes. **(d)** HCT intersection p-values for selected epigenetic writers within cold challenge-induced genes are indicated in the form of a heatmap. HCT intersection analysis was carried out as described in the Methods section. White cells represent p > 0.05 intersections. The intensity of the color scheme is proportional to the confidence of the intersection between HCTs for a particular node and genes induced (red, ↑) or repressed (blue, ↓) in each cold challenge contrast. Lower confidence (higher p) intersections are towards the yellow end of the spectrum and higher confidence (lower p) intersections are towards the brick red end of the spectrum. Full numerical data are in Supplemental File 1. TN: thermoneutral; RT: room temperature; SC: severe cold.

We next questioned whether the expression of epigenetic writers in BAT is altered in the response to different housing temperatures. We observe robust (FDR < 0.05) footprints within cold challenge-induced genes for numerous enzymes in the methyltransferase, demethylase, acetyltransferase and deacetylase classes (**Figure 1D)**. These include nodes with characterized or inferred roles in thermal regulation, such as the acetyltransferases *Crebbp*/CBP, EP300/p300, and *Kat2a*/*Gcn5*; deacetylases such as members of the HDAC and sirtuin families; methyltransferase members of the KMT2 family as well as *Mettl3*; and the KDM family demethylases (Artsi et al., 2019; Chen et al., 2021; Dovey et al., 2010; Emmett et al., 2017; Mutlu and Puigserver, 2021; Namwanje et al., 2019; Sambeat et al., 2017; Wang et al., 2020; Yan et al., 2021; Yao et al., 2017). We observe particularly strong footprints among cold challenge-induced genes for *Setd1a*, which has only recently been implicated in thermogenesis (Hoshii et al., 2022). In addition to these nodes, our analysis hints at roles for a number of nodes with no previously resolved role in the response to cold challenge, such as *Cdyl*, *Kat5*/TIP60, *Kat8*, *Ash2l*, *Ehmt2* and *Setdb1*.

### Establishing a mouse model to interrogate the BAT epigenetic cold response

To investigate epigenetic changes in BAT due to chronic thermogenic activation in response to mild or severe cold housing temperatures, tissues were collected from 10-week-old mice housed under thermoneutral (TN, 28°C), room temperature (RT, 22°C), or, for the last two weeks of the study, severe cold (SC, 8°C) conditions. Our data support prior reports that lower housing temperatures are associated with significant reductions in adiposity, as we observe decreased end body weight, total fat mass, and weight of gonadal white adipose tissue in SC- vs RT- or TN-housed mice (Zhao et al., 2022). There was no significant effect on nonfasting blood glucose levels, lean mass, or weights of BAT, liver, or spleen in our study (**Table S1**). Consistent with previous reports in hamsters and rats, kidney weight was inversely associated with housing temperature (Chaffee et al., 1963; Yahata and Kuroshima, 1989).

### DNA methylation and gene expression in BAT is altered in response to chronic thermogenic activation

DNA methylation at the promoters and first intron of genes is inversely correlated with gene expression, whereas intragenic DNA methylation may reduce spurious transcription and regulate the expression of alternate splice variants (Anastasiadi et al., 2018; Klose and Bird, 2006; Maunakea et al., 2013; Neri et al., 2017). To date, it is not known whether adaptation to cold induces large-scale epigenetic changes to DNA that impact global expression of genes and their splice variants in BAT. Reanalysis of the RNA-Seq data yielded numerous differentially expressed alternative splicing factors and alternative splice variants for BAT taken from SC- vs TN-housed mice (**Table 2**, **Figure 2A**, and **Supplemental Files S2 and S3**). To assess whether epigenetic modifications of DNA were occurring in BAT in response to chronic thermogenic activation, we first conducted reduced representation bisulfite sequencing (RRBS) using DNA extracted from BAT samples taken from mice housed under TN, RT, and SC temperature conditions. We noted 111 differentially hypomethylated and 18 differentially hypermethylated promoters; and 92 differentially hypomethylated and 16 differentially hypermethylated intragenic regions in BAT from RT-vs TN-housed mice (**Figure 2B-D**). Housing at SC further reduced promoter and intragenic region methylation when compared with TN. There were fewer differentially methylated promoters and intragenic regions in BAT taken from mice housed under SC vs RT conditions. These data show that changes in DNA methylation in BAT are associated with chronic activation of the NST response and that the more severe the chronic cold challenge, the greater the changes to DNA methylation.

**Figure 2.**
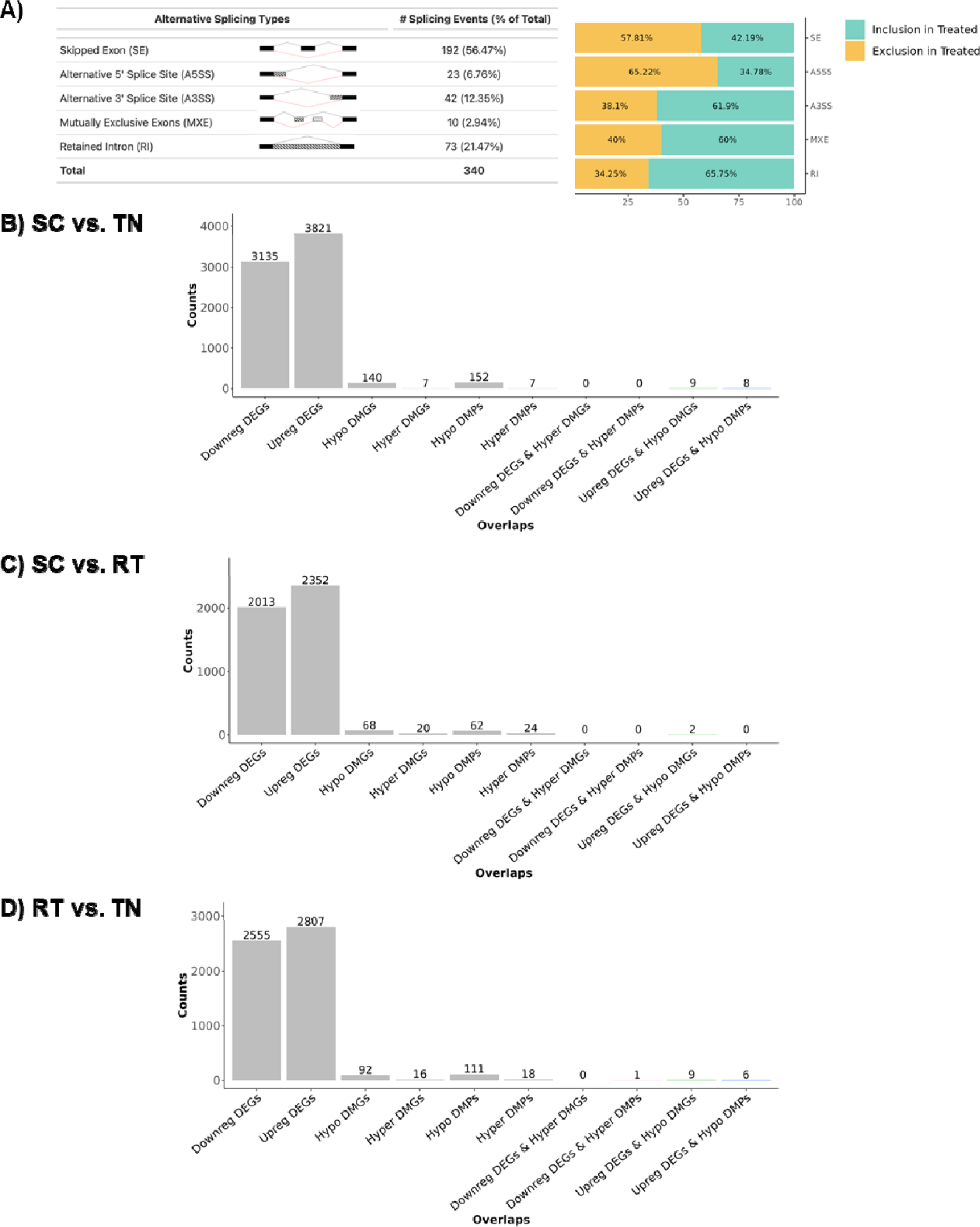
Alternative splice variants and reduced representation bisulfite sequencing (RRBS) data reveal changes in brown adipose tissue associated with housing temperature. **(a)** A comparison of SC vs TN housing RNA-seq data reveals differential expression of alternate splice variants in BAT. Integrated RNA-seq and RRBS data for **(b)** SC vs TN housed mice. **(c)** SC vs RT housed mice; and **(d)** RT vs TN housed mice. RRBS data shown are for n=4 per group with percent methylation difference > |5|, *p_adj_* < 0.05. DEG: differentially expressed genes. DMP: differentially methylated promoters. DMG: differentially methylated genes. TN: thermoneutral; RT: room temperature; SC: severe cold.

We next sought to identify genes with changes in promoter methylation that are inversely associated with gene expression, suggesting gene expression regulation resulting from altered DNA methylation. We integrated our RRBS and RNA-Seq data from C57BL/6J wildtype mice housed under similar conditions (**Table 3 and Supplemental File S3**) (Sanchez-Gurmaches et al., 2018). In mice housed at RT vs TN, *Car8*, *Ndufb11*, *Vma21*, *Vldlr*, *Lyplal1*, and *Car13 showed* the greatest percent reduction in promotor methylation that was also associated with increased gene expression in BAT (percent methylation difference > |5|; 1.5-1.8-fold expression; *p_adj_* <0.05). Conversely, increased promoter methylation for *Tpcn2* was associated with decreased expression (0.54-fold, percent methylation difference > |5|; 0.54-fold expression; *p_adj_* <0.05). SC housing resulted in hypomethylation of additional promoters, including the promoters for *Dlg3*, *Uba1*, *Dnajc14*, and *Tspan31*, for which gene expression was also significantly increased (1.15-2.65-fold, percent methylation difference > |5|; 1.15-2.65-fold expression; *p_adj_* <0.05). Although we observed significant differential methylation of promoters in BAT from SC- vs RT-housed mice, none were inversely associated with gene expression changes. Taken together, our findings from integrated RNA-seq and RRBS datasets suggest that differential methylation of DNA at promoters may play a role in the differential expression of genes in response to chronic severe cold in BAT.

**Table 3.**
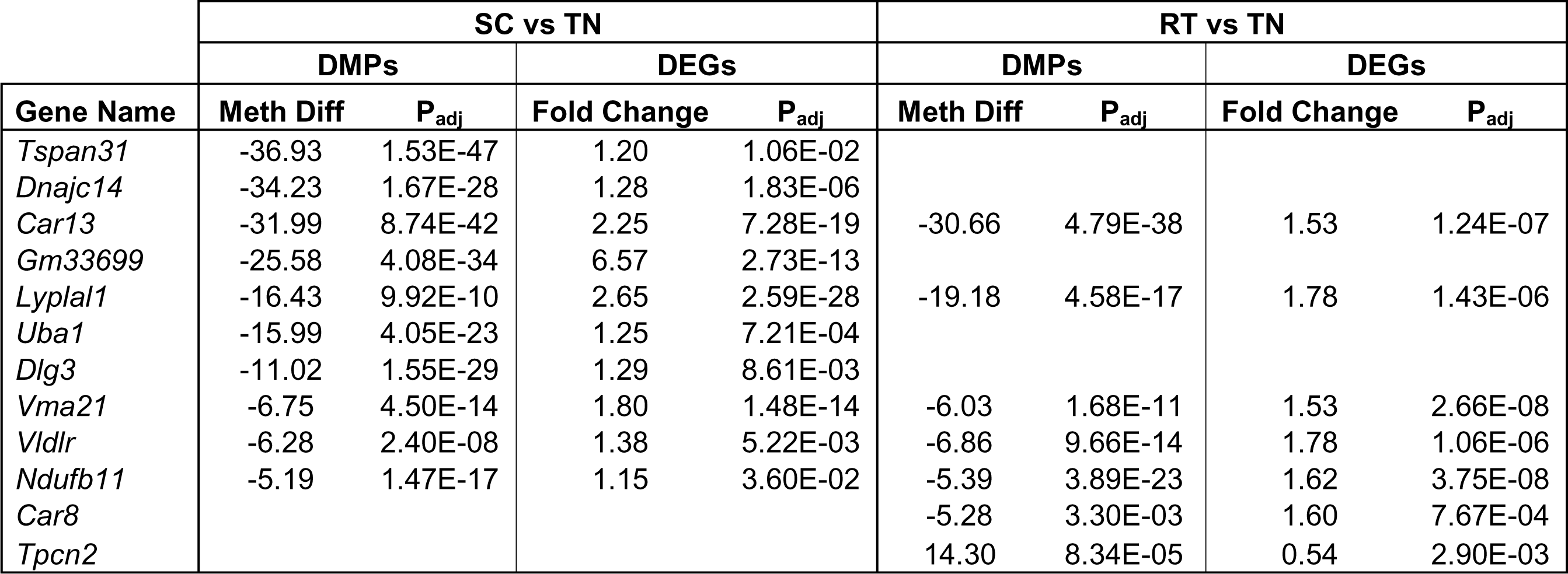
Integration of RRBS data with RNA-seq reveals genes that may be regulated by altered DNA methylation in response to different housing temperatures. DMPs: differentially methylated promoters; DEGs: differentially expressed genes; TN: thermoneutral; RT: room temperature; SC: severe cold.

### Quantitative, proteoform-level data can be obtained from brown adipose tissue

The capacity to rigorously and sensitively quantitate a wide dynamic range of histone proteoforms has only recently been established (Holt et al., 2019; Wang et al., 2018a). Furthermore, top-down mass spectrometry-based analysis of histone proteoforms has not been previously achieved for BAT in any study to date due to the high lipid content of the tissue. A “proteoform” is a protein defined with chemical precision, including how combinations of PTMs co-occur in cis, on single molecules (HThe Consortium for Top Down Proteomics et al., 2013). Thus, quantitation of histone proteoforms accesses the true physiological state and simultaneously provides quantitation of associated discrete attributes such as histone PTMs. To analyze changes in histone modifications in BAT due to different housing temperatures, we first developed histone extraction methods for brown adipose tissue, building on our protocol for histone extraction from cells (Holt et al., 2021). Considering the high lipid content of BAT, we performed a matrix of experiments changing the percent detergent (NP-40, 0.3-1%), the number of times incubated with detergent, the incubation time in detergent, the number of subsequent washes of nuclei, and the duration of centrifugation. Successful changes that were incorporated in this protocol include an additional incubation with detergent, an additional wash of nuclei, and a longer centrifugation duration compared to our previously published protocol (Holt et al., 2021). A summary of histone yield with chosen experiments is shown in **Table S2**. Acid extraction was successful with no changes to the protocol. Once histones were isolated, offline HPLC was used to separate histone families and H3 variants and our published mass spectrometry methods could be used (Holt et al., 2021, 2019, p. 4). Our final workflow is shown in **Figure 3A**. We obtained adequate offline HPLC separation of histone family members yielding sufficient quantity for LC-MS/MS analysis (**Figure 3B, Table S3A**). This allows for very efficient histone extraction, yielding more histone per gram tissue than obtained with livers (**Figure 3C** and **Table S3B**). Once these histones were prepared and analyzed by mass spectrometry, we obtained high-quality chromatography, localization of PTMs, and quantitation of proteoforms (**Figure 3D-G**). Overall, our methodological advancements enable histone isolation from brown adipose tissue for mass spectrometry analysis. Combined with existing methods, we obtain unbiased histone proteoform quantitation from BAT for the first time.

**Figure 3.**
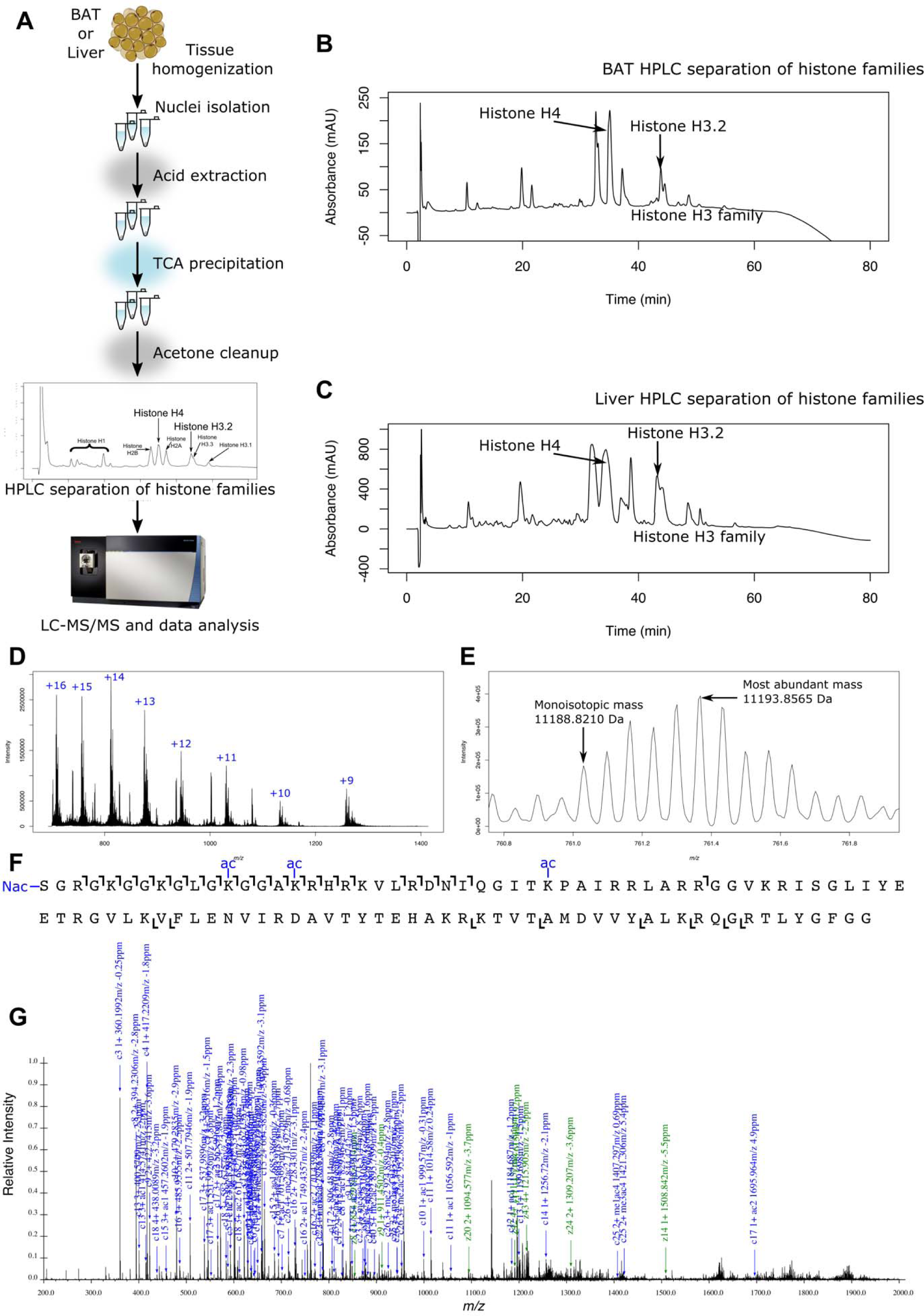
Workflow to obtain histone proteoform data from brown adipose tissue and liver. **(a)** The general workflow includes tissue homogenization, nuclei isolation, acid extraction, HPLC separation of histone families, and LC-MS/MS data acquisition. HPLC separation of histones shows a clean separation of histone families and H3 variants for **(b)** BAT and **(c)** liver. **(d)** MS1 average spectra of histone H4 extracted from BAT from 40 to 80 minutes show clear +16 to +9 charge states matching the *m/z* of histone H4. **(e)** The MS1 spectrum of 11,193.8565 Da species (761.2961 *m/z*, charge +15) matches a mass of H4 with N-terminal acetylation + 0 methylations + 3 acetylations + 0 phosphorylations with less than 10 ppm error. **(f)** The ion map from MS2 fragmentation of species from (e) shows unambiguous localization information for the proteoform H4 <N-acK12acK16acK31ac> from a mass change at less than 10 ppm error. **(g)** Annotated MS2 spectrum shows most abundant peaks are from the same proteoform, <N-acK12acK16acK31ac>, with less than 10 ppm error. Together, these methods enable the absolute quantitation of BAT and liver histone proteoforms.

### Quantitative analysis of histone post-translational modifications and proteoforms

Proteoforms are chemically defined single protein molecules, complete with all sources of variation (The Consortium for Top Down Proteomics et al., 2013). This includes how PTMs co-occur, in cis, on the same molecule and on which sequence variant (Dang et al., 2016; Joseph and Young, 2023). Thus, quantitative proteoform-level data presents a more complete description of the true physiological state of histone proteins (Taylor and Young, 2021). To distinguish PTMs and proteoforms we use curly brackets “{}” to indicate that the enclosed PTMs are present, and the abundance of all proteoforms that contain those modifications are summed. Angle brackets “<>” indicate that the enclosed PTMs are the only modifications on the histone protein. For example, H3<K9ME2K14ACK23ACK27ME3> is one of the most common states of histone H3 (Young et al., 2009).

### Cold adaptation results in tissue-specific changes in total histone H3.2 and H4 acetylation in BAT, but not methylation

The observed housing temperature-dependent differences in the expression of genes involved in acetyl-CoA generation and one-carbon metabolism in BAT suggests that the pool of available acetyl-CoA and methyl donor (*S*-adenosyl methionine, SAM) for histone modification may also be influenced by the degree of thermogenic activation (**Table 2**). Therefore, we sought to determine the effect of housing temperature on bulk histone acetylation and methylation in BAT. To understand the unique basal epigenetic state of BAT, we also analyzed histones from liver samples taken from the same animals for comparison. The percent of histone with one or more acetylations changes for both histone H3.2 and H4 in BAT (**Figure 4A**). Histone H3.2 with at least one acetylation is more abundant in BAT at SC compared to TN (45.2 and 35.6 percent H3.2 that has at least one non-N terminal acetyl group). Histone H3.2 acetylation in BAT is more abundant at RT compared to TN (43.1 and 35.6 percent acetylated). Histone H4 acetylation is also the most abundant in BAT at SC compared to TN (39.4 and 36.9 percent acetylated, *p* = 0.03) and SC compared to RT (39.4 and 36.8 percent that is acetylated, *p* = 0.02). However, there is no significant effect of housing temperature on bulk histone acetylation in liver. There are also no significant differences between tissues with histone H3.2 or H4 acetylation at each housing temperature. The percent of histones with at least one methylation did not change significantly with housing temperature or tissue type (**Figure 4B**). As noted above, these high stoichiometric occupancies are expected because H3 is physiologically hyper-modified and we are considering multiple sites of methylation in bulk. Our analysis approach allows for absolute quantitation and more directly reflects the stoichiometric use of the acetyl-CoA pool, the number of acetyl or methyl groups per histone molecule, and is shown in **Figure 4C** and **4D** respectively. Histone H3.2 acetylation follows a similar trend where SC BAT shows the highest acetylation levels, 0.63 acetylations per histone H3.2 molecule, except for liver at RT which is 0.65 acetylations per H3.2. There are large differences in acetylations per histone H3.2 molecule with a 0.12 difference between SC and TN housing and 0.10 difference between RT and TN housing. Histone H4 acetylation is highest in BAT from mice housed at SC, 0.46 acetylations per histone H4 molecule, and second highest in liver from mice housed at SC, 0.45 acetylations per H4 molecule. Taken together, our data show for the first time that bulk changes to histone acetylation occur in BAT with cold adaptation and are tissue-specific.

**Figure 4.**
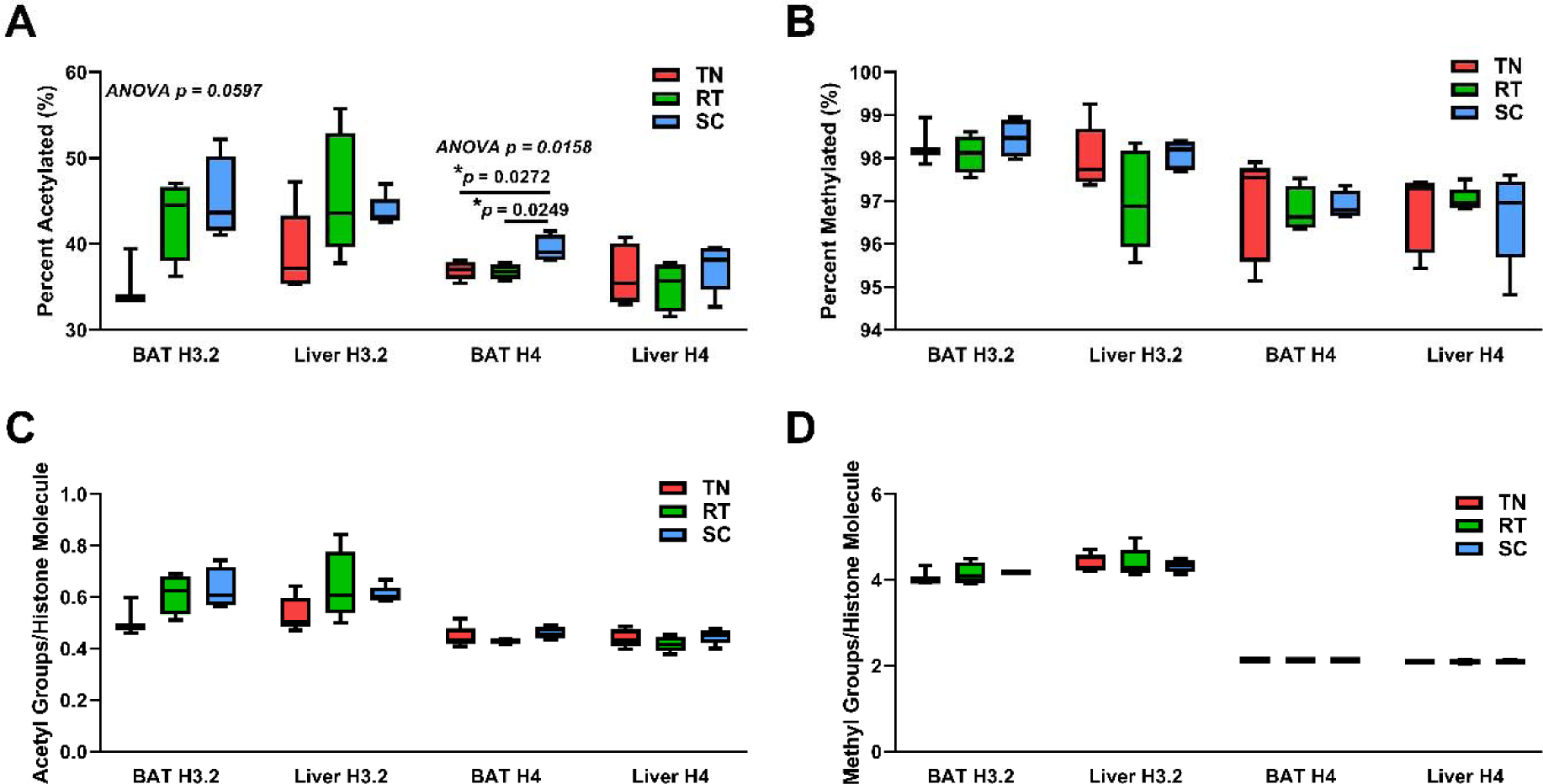
Housing temperature alters bulk histone acetylation, but not methylation, in BAT and has no effect on bulk histone acetylation or methylation in liver. Percent **(a)** acetylated (1+ acetylation) and **(b)** methylated (1+ methylation) histones H3.2 and H4 isolated from liver and BAT. Number of **(c)** acetyl groups and **(d)** methyl groups (K4me2 = 2 methyl groups) per histones H3.2 and H4 isolated from liver and BAT. One-way ANOVA testing was conducted for all histone data from each tissue comparing TN, RT, and SC housing conditions, and *p*-values are indicated in each plot. TN: thermoneutral; RT: room temperature; SC: severe cold.

### Brown adipose tissue exhibits different histone PTMs and proteoforms than liver

We next sought to determine whether histone PTMs and proteoforms are different in BAT versus liver. Here, we focus on significant differences at RT between tissues for discrete PTMs, binary PTM combinations, and proteoforms (**Figure 5 and Supplemental File S4**). However, the highest number of significant differences between tissues is at SC. In general, K9me3 and K36un and their combinations are more abundant in BAT compared to liver. Exceptions include {K27ac,K36un} at RT and {K27me3,K36un} at SC. At RT, BAT histone H3.2 shows significantly higher levels of PTMs K9me1, K9me3, K27me1, and K36un. Many differences are also observed between tissues for binary modifications and proteoforms in histone H3.2 (**Figure 5A** and **5D**). Because of our absolute quantitation, we can calculate the difference – not just fold change – in the abundance of PTMs or proteoforms. This difference is indicative of the percent of the genome that is affected by the change in this proteoform (percentage point, pp). For example, H3.2<K9ME2K27ME1> is 1.7 percent abundant in the liver and 6.0 percent abundant in BAT, a 4.3 pp difference. Thus, the difference in abundance of H3.2<K9ME2K27ME1> affects a massive 4.3 percent of the genome when comparing liver and BAT at RT. For reference, the abundance of {K4me3} typically decorates less than one percent of the genome yet is crucial for active gene transcription (Jain et al., 2023). The largest absolute differences with significance are observed with the histone H3.2 proteoforms <K9ME3K27ME1>, <K9ME2K27ME1>, and <K9ME2K27ME3K36ME1>. H3.2<K9ME3K27ME1> and <K9ME2K27ME1> are more abundant in BAT than in the liver. H3.2<K9ME3K27ME1> is 0.5 percent abundant in the liver and 3.3 percent abundant in BAT, affecting 2.7 percent of the genome. H3.2<K9ME2K27ME3K36ME1> is more abundant in the liver than BAT (3.7 and 1.6 percent of total H3.2, respectively) and affects 2.1 percent of the genome. In total, these proteoforms are responsible for a shift in 10.8 percent of the genome between BAT and liver.

**Figure 5.**
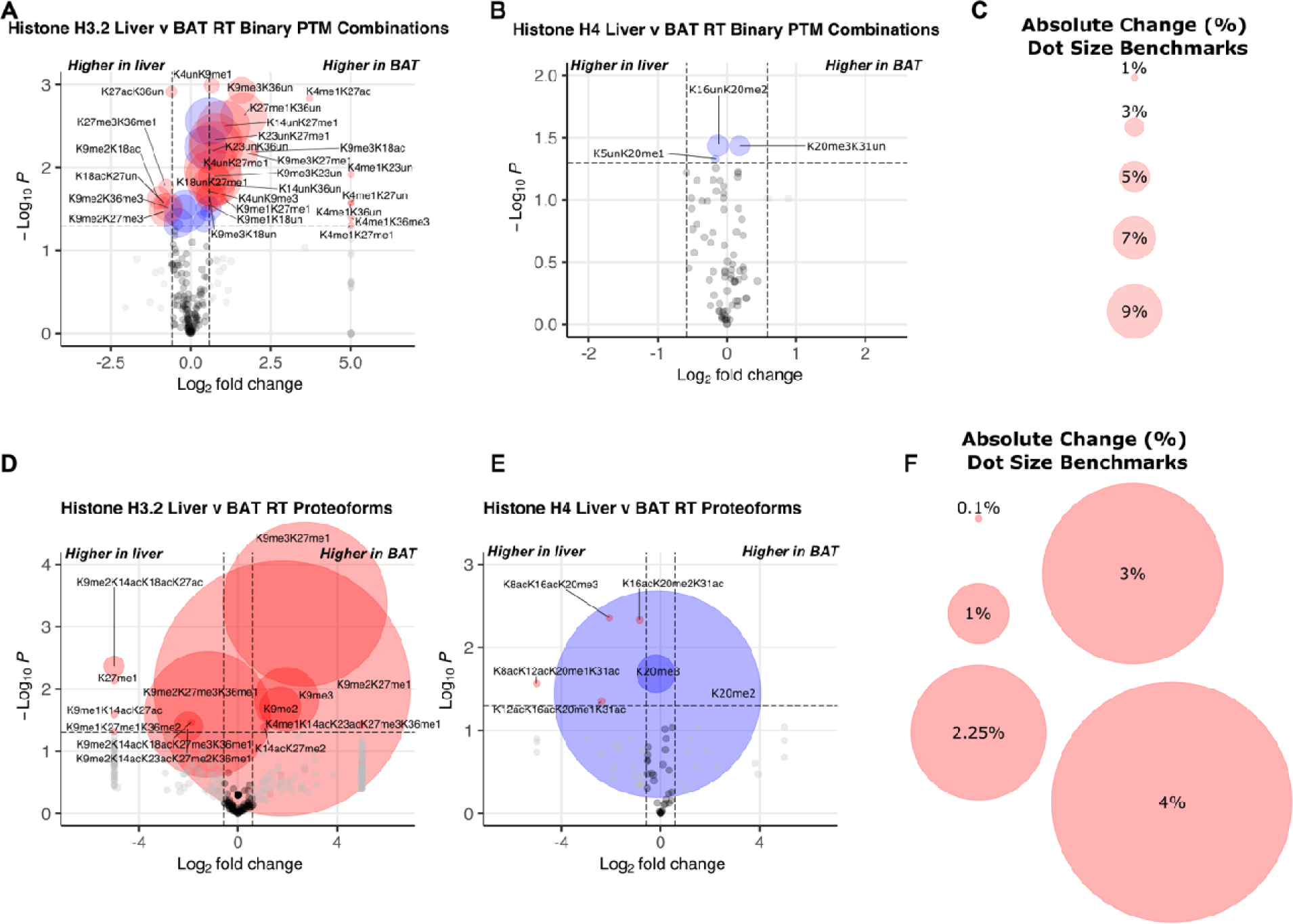
Under normal room temperature housing conditions, BAT histone H3.2 and H4 post-translational modifications and proteoforms are distinct from those observed in liver. **(a)** Histone H3.2 and **(b)** histone H4 binary PTM combinations in BAT and liver. **(c)** Dot size references for binary volcano plots (*a,b*). **(d)** Histone H3.2 and **(e)** histone H4 proteoforms from BAT and liver. **(f)** Dot size references for proteoform volcano plots (d.e). P-values were calculated using Welch’s t-test. Volcano plots show cutoffs of 1.5 for fold change and *p* < 0.05. Red dots represent both p < 0.05 and greater than 1.5-fold change; blue dots represent p < 0.05 and less than 1.5-fold change; grey dots represent p > 0.05 and greater than 1.5-fold change; black dots represent p > 0.05 and less than 1.5-fold change. RT: room temperature.

Overall, histone H4 PTM changes are observed most frequently at SC (**Supplemental File S4**). This includes {K20me1} containing binary combinations and the otherwise unmodified proteoform <K20ME1> which are more abundant in the liver compared to BAT. Histone H4 at RT shows no significant differences between tissues for discrete PTMs or binary PTM combinations, proteoform abundances (**Figure 5B** and **5E**) trend toward differences in BAT and liver. H4<K20ME1> and <K20ME2> are more abundant in liver than BAT. H4<K20ME1> is 5.4 percent abundant in the liver and 4.7 percent abundant in BAT, changing 0.62 percent of the genome. H4<K20ME2> is 39.4 percent abundant in the liver and 36 percent abundant in BAT, causing a shift in 3.4 percent of the genome while only 0.91-fold change. H4<K20ME3> is more abundant in BAT than liver. H4<K20ME3> is 19 percent abundant in the liver and 22 percent abundant in BAT, causing a shift in 2.5 percent of the genome. Overall, our results reveal tissue-specific discrete PTM abundances, PTM combinations, and proteoforms for histone H3.2, and to a lesser extent, H4 at RT housing.

### BAT histone H3.2 changes with cold exposure are centered around K9me2 and K9un

Few temperature-dependent changes in histone H3.2 PTMs and proteoforms were observed in liver (**Supplemental File S5**). Our motivating central hypothesis is that there is a BAT-specific epigenetic response to changes in housing temperature. This is rationalized by the importance of histone PTMs in the regulation of transcription and the known unique physiological role of BAT in nonshivering thermogenesis. To determine epigenetic changes that occur during the cold response, we analyzed discrete PTMs, PTM combinations, and proteoforms of histones in BAT from TN-, RT-, and SC-housed mice. Discrete PTM analysis of histone H3.2 shows significant increases in PTMs K9me2 and combined K9me2 and K9me3, {K9me2/3}; and decreased unmodified K9, {K9un}, in response to SC housing (**Figure 6A**). Histone H3.2 proteoforms show trends in K9 di-methylation with K36 methylation (**Figure 6B)**. The presence of K9me2 or K9me3 and K36me1 on the same molecule is greatest at SC when combined with K23ac (**Figure 6C**). Supplemental data is available for further analysis (MassIVE: MSV000092105, **Supplemental File S5**). Our data indicate a role for altered histone H3.2 methylation and acetylation in the regulation of transcription in BAT in response to cold stress.

**Figure 6.**
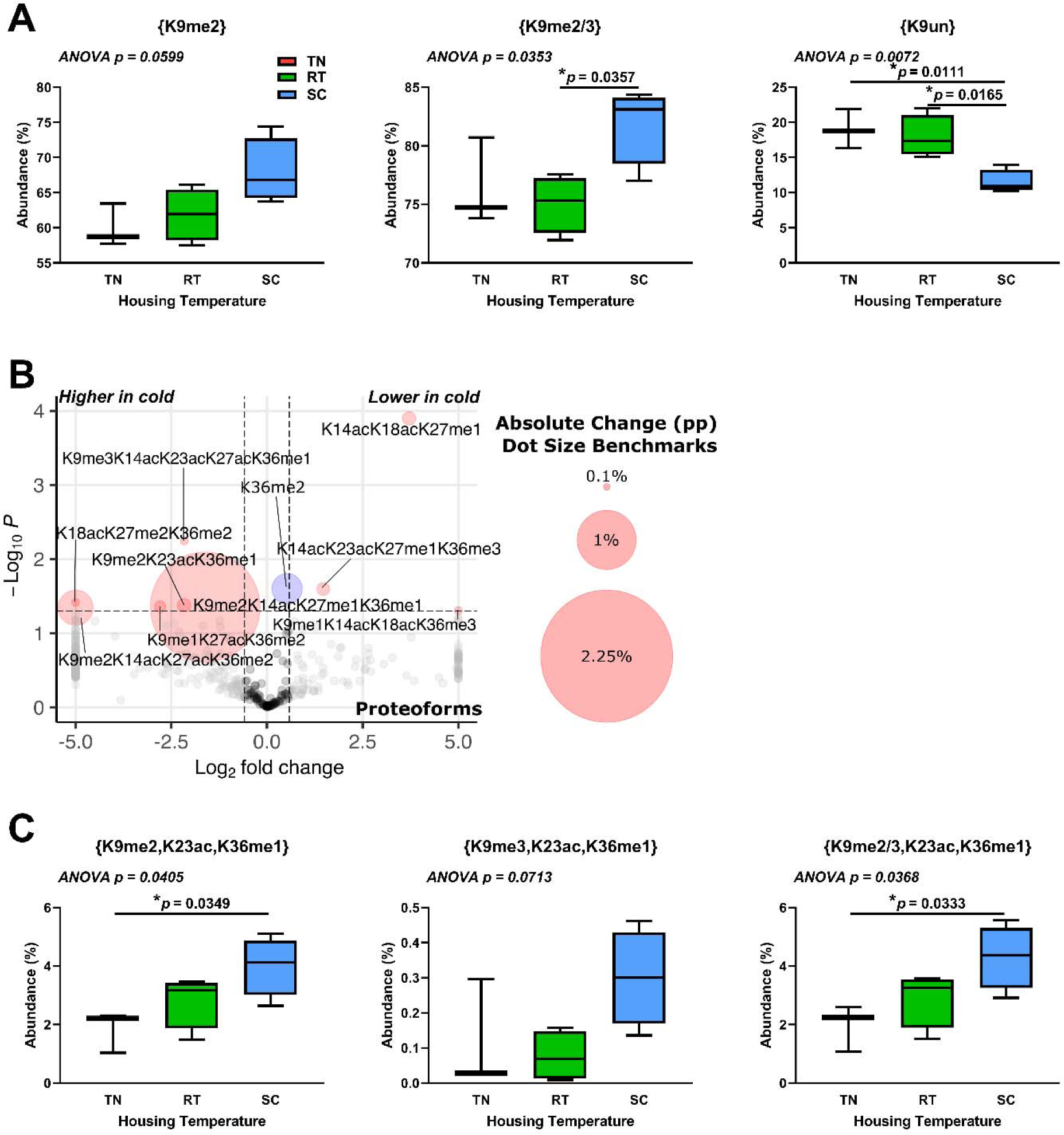
Histone H3.2 K9me2- or K9me3-containing proteoforms increase in brown adipose tissue in response to cold housing. (a) Selected discrete H3.2 PTM abundance at different housing temperatures. **(b)** Proteoform changes between SC and TN. Dot size corresponds to absolute percentage point change. Red dots represent both p < 0.05 and greater than 1.5-fold change; blue dots represent p < 0.05 and less than 1.5- fold change; grey dots represent p > 0.05 and greater than 1.5-fold change; black dots represent p > 0.05 and less than 1.5-fold change. **(c)** Ternary (3-PTM) combinations show significant differences with K9 di- and tri-methylation, K23 acetylation, and K36 monomethylation. Welch’s t-test was used for volcano plot data, with cutoffs of 1.5 for fold change and *p* < 0.05. One-way ANOVA testing was conducted for all PTM data comparing TN, RT, and SC housing conditions, and *p*-values are indicated in each plot. TN: thermoneutral; RT: room temperature; SC: severe cold.

### Histone H4 proteoforms in cold-adapted brown adipose tissue show highly specific changes including K16ac

We hypothesize that there are BAT-specific epigenetic changes with cold exposure that will affect histone H4 proteoforms. As with histone H3.2, we analyzed discrete PTMs, PTM combinations, and proteoforms for histone H4. We observe no significant histone H4 discrete PTM changes between BAT from mice housed at various temperatures. Interestingly, most H4 proteoforms that significantly change with housing temperature include K16ac, which may play a permissive role in the regulation of gene expression (**Figure 7A-B**) (Hilfiker et al., 1997; Suka et al., 2002). Further examination reveals specific proteoform changes, which includes <K12ACK16ACK20ME1K31AC> and <K8ACK16ACK20ME3>, both of which significantly increase at SC vs RT or SC vs TN (**Figure 7C** and **7D**). Supplemental data is available for further analysis (MassIVE: MSV000092105, **Supplemental File S6**). These results show that specific histone H4 proteoforms increase in BAT in response to chronic cold.

**Figure 7.**
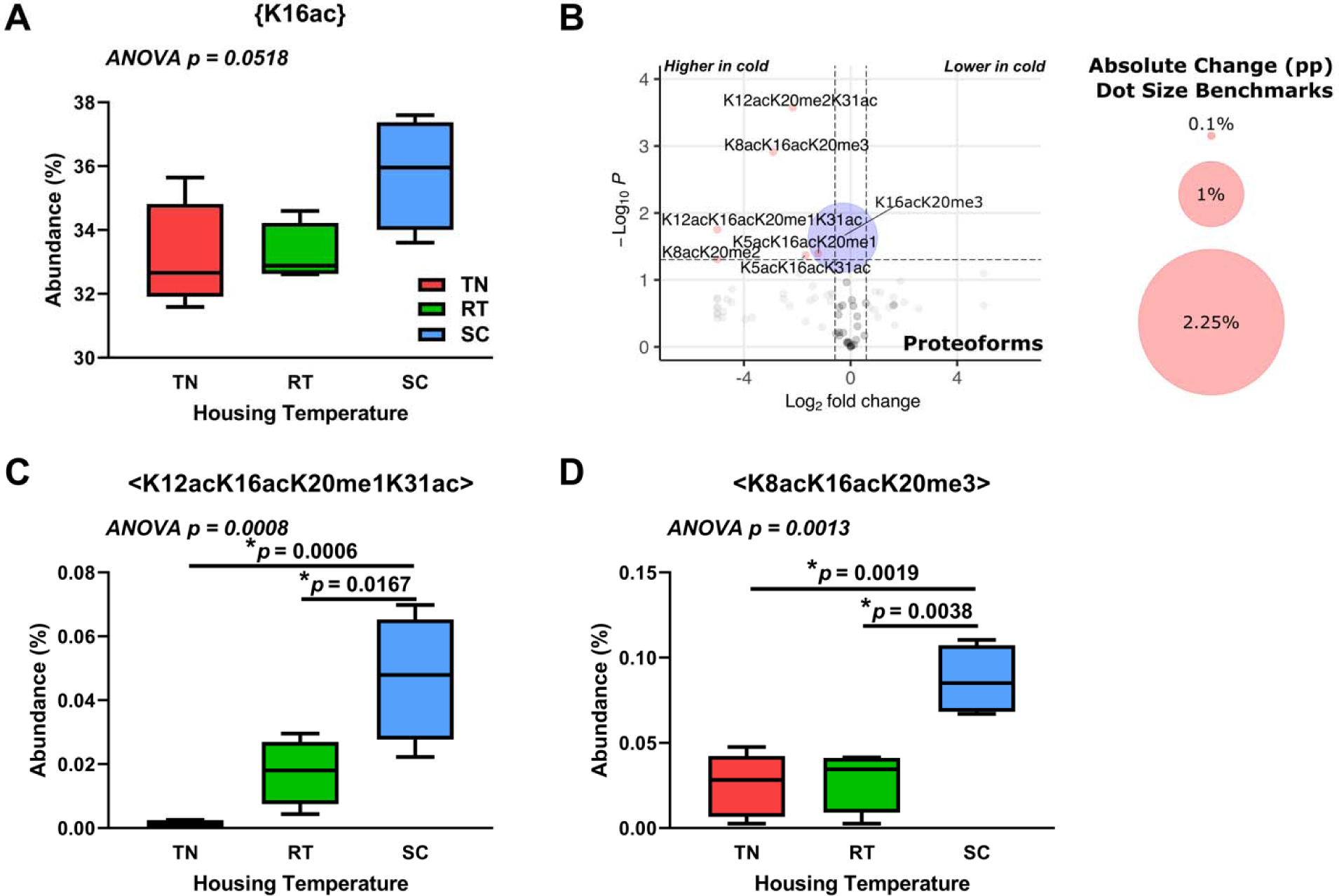
Housing temperature significantly alters histone H4 K16ac-containing proteoforms in brown adipose tissue. (a) Discrete K16ac PTM marks on histone H4 at different housing temperatures. **(b)** Histone H4 proteoforms from BAT from SC vs TN housing temperatures. Dot size corresponds to absolute percentage point change. Red dots represent both p < 0.05 and greater than 1.5-fold change; blue dots represent p < 0.05 and less than 1.5- fold change; grey dots represent p > 0.05 and greater than 1.5-fold change; black dots represent p > 0.05 and less than 1.5-fold change. **(c)** H4<K12ACK16ACK20ME1K31AC> proteoform abundance at different housing temperatures. **(d)** H4<K8ACK16ACK20ME3> proteoform abundance at different housing temperatures. Welch’s t-test was used for volcano plot data, with cutoffs of 1.5 for fold change and *p* < 0.05. One-way ANOVA testing was conducted for all data comparing TN, RT, and SC housing conditions, and *p*-values are indicated in each plot. TN: thermoneutral; RT: room temperature; SC: severe cold.

## Discussion

The current work presents a novel method for quantitative top-down proteomic analyses of histone modifications in brown adipose tissue, which we cross-validate with a multi-omics approach. This method enables us to carry out the first-ever in-depth exploration of changes in histone modifications in BAT due to differing degrees of thermogenic activation. Our proteomic results are supported by analyses of gene expression that indicate changes in epigenetic modifiers, as well as DNA methylation changes, showing broad epigenetic changes occur as a result of chronic thermogenic activation. We observe significant temperature-dependent changes in the expression of genes that regulate one-carbon metabolism, DNA methylation, histone modification, and alternative splicing. We further uncover a number of alternative splice variants that are differentially expressed in response to thermogenic activation. We note decreased promoter and intragenic DNA methylation with colder housing temperatures, and this is associated with reduced *Dnmt1*, *Dnmt3a*, and *Dnmt3b* expression. Consistent with a functional effect of increased expression of acetyl-CoA generating enzymes in BAT, we observe tissue-specific increases in H3.2 and H4 total acetylation in response to severe chronic cold. Further examination reveals tissue-specific changes in histone H3.2 and H4 PTMs and proteoforms in BAT in response to chronic thermogenic activation. Our data show for the first time that chronic thermogenic activation results in large-scale epigenetic changes to both DNA and histones in murine BAT that represent both transcriptional “on” and “off” signals. These signals indicate an important role in epigenetic regulation of BAT adaptation to chronic thermogenic activation.

### Housing Temperature Alters DNA Methylation in BAT

We report the first analysis of global thermogenesis-associated DNA methylation changes in BAT. Using RRBS, we discovered reduced DNA methylation in BAT as housing temperature decreases. This may result from reductions in DNA methyltransferase activity, as RNA-seq reveals lower expression of *Dnmt1*, *Dnmt3a*, and *Dnmt3b* at both SC and RT compared to TN (Anastasiadi et al., 2018; Klose and Bird, 2006; Li et al., 2021; Tovy et al., 2022; Wang et al., 2021). The maintenance DNA methyltransferase DNMT1 represses myogenic remodeling of BAT, and inhibits the expression of adiponectin (*Adipoq*) by adipocytes (Kim et al., 2015; Li et al., 2021). Additionally, global inhibition of DNMT1 in diabetic obese mice (db/db) *in vivo* improves whole-body glucose homeostasis in an adiponectin-dependent manner. However, expression of adiponectin, which has been shown to inhibit BAT thermogenesis, is significantly reduced by SC compared with RT or TN housing conditions (Qiao et al., 2014; Sanchez-Gurmaches et al., 2018). We also found no association between *Adipoq* expression and DNA methylation at its promoter, suggesting that DNMT1 activity does not play a major role in the regulation of *Adipoq* expression in BAT during chronic severe cold stress. BAT-specific *Dnmt1*-deletion has no effect on body weight or glucose homeostasis when mice are fed a standard low fat diet (Park et al., 2021). However, thermogenic response to severe cold was not tested in these mice. Both DNMT3a and DNMT3b regulate *de novo* methylation of DNA and play important roles in adipocyte development and differentiation. DNMT3a regulates preadipocyte differentiation (Tovy et al., 2022). DNMT3b deficiency in Myf5^+^ brown adipocyte-skeletal muscle precursor cells reduces thermogenic gene expression and upregulates myogenic gene expression in BAT of female mice (Wang et al., 2021). Further studies are warranted with regard to the roles of DNA methyltransferases in adult BAT in response to chronic cold.

We also report novel findings from integration of our RRBS data with RNA-seq data. Interestingly, the genes with the greatest reductions in promoter methylation for which expression is also increased in response to chronic cold stress include *Vldlr*, recently reported to play an important role in fatty acid uptake by thermogenically active BAT, and *Ndufb11*, a component of mitochondrial complex I (Carroll et al., 2002; Shin et al., 2022). Of note, BAT *Vldlr* expression and uptake of VLDL increase in response to cold stress, and VLDLR deficiency impairs thermogenesis in mice (Shin et al., 2022). As VLDLR also facilitates the uptake of fatty acids from chylomicrons via lipoprotein lipase-mediated triglyceride hydrolysis, increased *Vldlr* expression represents a means through which cold stress may promote the uptake of fatty acids from circulating chylomicrons postprandially, as well (Goudriaan et al., 2004). Our results suggest that alterations in DNA methylation at promoters is one mechanism through which gene expression for optimal nutrient utilization during thermogenesis is regulated during BAT adaptation to prolonged cold stress.

Methylation of intragenic regions of DNA has been shown to differentially affect the expression of alternative splice variants and modification of histones (Lorincz et al., 2004; Maunakea et al., 2013). Our RNA-seq analysis yields a number of alternative splice variants that are differentially expressed in BAT at SC vs TN. Additionally, we note associated increased expression of HuR/*Elavl1* and *Igf2bp2*, and decreased expression *Nova2*, *Pspc1*, and many serine-arginine rich splicing factors in BAT from mice housed at SC vs TN. This suggests that expression of RNA-binding proteins known to mediate alternative splicing in adipose tissues is also altered during chronic cold stress (Chao et al., 2021; Vernia et al., 2016; Zhang et al., 2022). Our results indicate the need for further studies to better characterize the mechanism(s) driving alternative splicing events in BAT, as well as the biological function of alternative splice variants that are expressed in BAT, in response to chronic cold stress.

Here, using our newly developed method of top-down mass spectrometry-based analysis of histone PTMs and proteoforms in BAT, we also describe a number of histone modifications, particularly increased bulk acetylation, H3.2{K9me2}, and H3.2{K9me3}. DNA methylation has been shown to facilitate the recruitment and methylation of H3K9, and loss of DNMT1 activity is associated with increased acetylation and decreased H3K9 di- and trimethylation, (Espada et al., 2004, p. 1; Fujita et al., 2003; Fuks et al., 2003). Additionally, H3K9me3 reinforces DNMT1 activity and genome targeting (Ren et al., 2020). This suggests that decreased *Dnmt1* expression may play a role in the increased histone acetylation we observe, and perhaps increased H3.2{K9me2} or {K9me3} ensures DNMT1 fidelity with decreased expression. Overall, our results show that chronic cold stress affects DNA methylation in BAT, and that this may play a role in adaptation through effects on transcription, transcript splicing, and histone modifications. RRBS can only assess methylation at 10-15% of all CpG sites in the genome and that cannot distinguish between 5-methylcytosine and its demethylation intermediate, 5-hydroxymethylcytosine (Gu et al., 2011; Huang et al., 2010; Jin et al., 2010). Future studies will utilize whole genome bisulfite sequencing for a more comprehensive approach to thermogenically-responsive changes in DNA methylation in BAT.

### Our Data Reveal the First Top-down Mass Spectrometry-based Quantitative Look at Histone Modifications in Murine BAT

The work described here is the first basal quantitative description and unbiased quantitative statistical analysis of changes in BAT histone PTMs and proteoforms. Top- or middle- down mass spectrometry-based methods are ideal for the unbiased identification and quantitation of histone PTMs (Han et al., 2006; Moradian et al., 2014; Pandeswari and Sabareesh, 2019; Patrie, 2016; Taylor and Young, 2021; Tran et al., 2011; Tvardovskiy et al., 2015). PTMs on histones are well-known to affect the binding of downstream factors that effectuate genome function (Plazas-Mayorca et al., 2010; Vaquero et al., 2007; Wang et al., 2018a). As described in our recent review, histone modifications often function in concert on single molecules (Taylor and Young, 2021). Isolation of nuclei from tissue presents many challenges due to the complexity of BAT tissue. We have recently published methods to overcome issues in most types of tissue (Holt et al., 2021; Taylor and Young, 2023). Here we optimize BAT nuclei isolation for subsequent acid extraction of histones and analysis of histone proteoforms by mass spectrometry (Holt et al., 2019, p. 4). A proteoform biology approach brings valuable new insights, such as how multiple PTMs function in concert on single molecules to affect BAT function, while also providing quantitation of more conventional discrete PTMs. Our combined set of transcriptional footprint, splice variant, DNA methylation, and histone PTM and proteoform analyses serves as a valuable resource for future interrogations of the epigenetic response of brown adipose tissue due to chronic thermogenic activation.

### BAT and Liver are Epigenetically Distinct

Our proteomic analysis of histones H3.2 and H4 in BAT and liver from mice housed at different temperatures reveals tissue-specific responses to cold. Histones in BAT are more acetylated at SC compared to other housing temperatures. In liver, there is no significant effect of housing temperature on histone acetylation levels. Additionally, there is no difference in bulk histone acetylation levels between tissues. Although our focus for comparison of proteoforms in these tissues was RT housing temperature, we observe the greatest number of significant differences in single PTMs, 2-PTM combinations, and proteoforms between tissues at SC. The significantly differing combinations containing H3.2 are always more abundant in BAT compared to liver, indicating transcriptional repression of genes through HP1 in BAT (Bannister et al., 2001). The abundance of H3.2{K36un} and combinations including K36un are typically more abundant in BAT compared to liver, consistent with decreased deposition of K36 methylation during transcription (Morris et al., 2005). Thus, there is more K9 methylation to recruit HP1, which would result in less transcription and less co-transcriptional methylation of K36. Fewer differences between tissues are observed with histone H4 but center on {K20me1} at SC. H4{K20me1} binary combinations and H4<K20ME1> are more abundant in the liver. Generally, we observe PTM differences between tissues that indicate less active transcription in BAT than in liver, and these differences are most pronounced under severe chronic cold housing conditions.

At RT, histone H3.2 proteoforms that are significantly more abundant in BAT than liver are H3.2<K9ME2K27ME1>, H3.2<K9ME3K27ME1>, H3.2<K9ME3>, and H3.2<K9ME2> and indicate an increase in heterochromatin in BAT (Boros et al., 2014; Vaquero et al., 2007). Histone H4 proteoforms H4<K20ME3>, H4<K16ACK20ME2>, and H4<K16ACK20ME3> trend toward greater abundance in BAT compared to liver. These proteoforms indicate transcriptional repression or a poised state (Schotta et al., 2004). H4<K20ME2> and H4<K20ME1> are significantly more abundant in liver than BAT and are associated with transcriptional activation (Wang et al., 2008). H4<K20ME2> is 39 percent of all H4 in liver and 36 percent abundant in BAT, affecting 3.4 percent of the genome. H4<K20ME1> is 5.4 percent of all histone H4 in liver and 4.7 percent abundant in BAT. Significant discrete changes between tissues are observed in histone H3.2 but not histone H4 at RT. The changes in proteoform abundances reflect poised positions that act as a scaffold for other factors that determine downstream effects (Wang et al., 2018a). These proteoforms are often quite abundant in tissues and cultured cells and their shift indicates a broad difference between tissue epigenetic regulation (Holt et al., 2019; Wang et al., 2018a). Significantly changing proteoforms of histone H3.2 and H4 indicate higher levels of heterochromatin in BAT compared to liver at RT (Boros et al., 2014; Schotta et al., 2004; Vaquero et al., 2007; Wang et al., 2008). Our histone proteoform findings also suggest that fewer genes are being actively transcribed in BAT under this condition of mild chronic cold activation.

### Housing Temperature Affects Histone Modifications in BAT

Our data show that BAT responds epigenetically to cold stimuli and further reveal changes in its epigenome that may drive its function during cold adaptation. Bulk histone acetylation increases in BAT from SC- vs TN-housed mice. This is consistent with changes in gene expression upon cold challenge. Expression of *Acss2* and *Chrebp (Mlxipl)* and its known target *Acly* increase in BAT when mice are housed under chronic mild and severe cold conditions (Iizuka et al., 2004; Sanchez-Gurmaches et al., 2018). ACLY and ACSS2 provide acetyl-CoA for histone acetylation through citrate and acetate, respectively, and their upregulation provides potential mechanisms for our findings. Additionally, carnitine acetyltransferase (CrAT) and carnitine octanoyltransferase (CrOT) have both been shown to play important roles in the ACLY- and ACSS2-independent conversion of short- and medium-chain acetylcarnitines to acetyl-CoA (Hsu et al., 2023; Izzo et al., 2023). Analysis of RNA-seq data reveals that both *Crat* and *Crot* expression increase in BAT during chronic cold housing, suggesting additional mechanisms through which BAT activation generates acetyl-CoA that can be used for post-translational modification of histones. We also observe a significant cold-induced transcriptional footprint of histone acetyltransferases CBP/p300 in BAT that is associated with the increased acetylation of histones H3.2 and H4. CBP/p300 are generally considered to be the most active histone acetyltransferases and play a crucial role in regulating adipocyte plasticity (Namwanje et al., 2019). CBP and p300 both preferentially acetylate histones H3 and H4 (Henry et al., 2013; Holt et al., 2021). Indeed, histone H4 and H3.2 acetylation sites are the most occupied at SC in BAT, indicating a shift in BAT acetyl-CoA pools. Taken together, our findings are consistent with increased intracellular acetyl-CoA levels and histone acetylation via p300/CBP contributing to increasing histone acetylation under conditions of chronic cold stress.

Specific histone post-translational modifications (PTMs) additionally suggest a mechanism through which temperature-dependent selective transcription in BAT may occur. Either the presence or absence of PTMs can be biological signals. Under chronic severe cold conditions, histone H3.2 {K9me2} and {K9me2/3} increase and {K9un} decreases, indicating an increase in repressive transcriptional signals (Boros et al., 2014; Vaquero et al., 2007). However, further investigation into the K9me2 mark reveal interplay with K36 methylation (K36me1 and K36me2) and such combinations enable K9me2 to affect transcriptional repression of some genes while others are transcribed (Morris et al., 2005). The decrease in {K9un} supports this model. While the lack of commercially available {K9un} antibodies means there are few studies implicating function, a reasonable interpretation of loss of {K9un} is a shift away from a more transcriptionally permissive but still poised state. {K9un} is directly prone to either acetylation (activation) or methylation (repression) without erasure of the antagonistic mark (Adamkova et al., 2017; Boros et al., 2014). As thermogenic activation increases, histone H4 proteoforms with K16ac increase. Importantly, {K16ac} acts as a scaffold and recruits factors for transcriptional activation (Taylor and Young, 2021). Mechanistically, H4{K20me2} promotes {K16ac} and further rapid acetylation in *cis* at K12, K8, and K5 (Wang et al., 2018a). In support of these PTM data, we note increased expression and transcriptional footprints of *Kat8* with thermogenic activation between SC- and RT- vs TN housing temperatures. KAT8 forms a specific H4K16 acetyltransferase complex with MSL1, MSL2, and MSL3 (Dou et al., 2005; Gupta et al., 2008; Smith et al., 2005). While a role for KAT8 in the thermogenic function of brown adipocytes has not been tested, KAT8 is required for adipocyte differentiation prior to clonal expansion in an *in vitro* culture model, (Burrell and Stephens, 2021). Taken together, histone H3 PTMs indicate persistent transcriptional repression of certain genes (through methylation) and histone H4 PTMs suggest an overall transient or poised state of transcription (through acetylation) in BAT when mice are housed at colder temperatures. These findings suggest a mechanism in BAT through which thermogenic activation fine-tunes transcription in a nutrient-sensitive manner via increased acetyl-CoA and KAT8 activity to modulate histone methylation-dependent gene repression.

Our gene expression and transcriptional footprint analyses suggests a regulatory mechanism of H3K9 methylation-dependent gene repression in BAT. RNA-seq analysis reveals that *Fcor* expression increases with decreasing housing temperature. FCOR regulates FOXO1 activity through repressive acetylation (Kodani and Nakae, 2020). We also observe an increased SIRT6 transcriptional footprint under colder housing temperatures. SIRT6 deacetylates FOXO1, and SIRT1 deacetylates SIRT6, thus increasing SIRT6 activity (Meng et al., 2020; Zhang et al., 2014). When not acetylated, FOXO1 forms a complex with SIRT1 and mediates the expression of SIRT1 (Kodani and Nakae, 2020; Xiong et al., 2011). Both SIRT1 and SIRT6 are known H3{K9ac} deacetylases (Kuang et al., 2018). Deacetylation of H3K9 allows for methylation (Adamkova et al., 2017). Also, SIRT1 mediated deacetylation of SUV39H1 at K266, activates this H3K9 dimethyltransferase, generating H3K9me3. (Vaquero et al., 2007, p. 1). The KDM6C footprint also increases upon cold challenge, which specifically demethylates H3K9me3 to K9me2 (Nic-Can et al., 2019). Thus, dramatically higher levels of {K9me2} at SC likely result from a transcriptionally repressive feedback loop with {K9me2/3} with cold challenge.

In summary, with the addition of data from our newly developed method for top-down mass spectrometry-based proteomic quantitation of histone proteoforms in BAT, we observe tissue-specific epigenetic responses to severe chronic cold stress. The transcriptional footprints of many histone modifying proteins, including both writers and erasers, increase upon cold challenge. The reduction in promoter DNA methylation and increased histone acetylation both suggest increased transcription, while increased histone H3.2 K9 di- and trimethylation indicates transcriptional repression under chronic cold conditions. These epigenetic changes, as well as alternative splicing, support an important role in adaptation to cold stress in BAT by regulating transcript expression. Future work will elucidate specific mechanisms of epigenetic regulation of BAT adaptation to chronic thermogenic activation.

## Materials and Methods

### Processing RNA sequencing data

Raw sequencing data was downloaded from Gene Expression Omnibus database under accession number GSE96681, using SRA (Sequence Read Archive) toolkit (Leinonen et al., 2011). Sequencing quality and adapter contamination were assessed using FastQC v0.11.9 (Andrews, 2010). Overall quality was determined to be satisfactory, and raw reads were aligned to the genome index using STAR v2.7.9a (Dobin et al., 2013). The STAR genome index was created using raw FASTA and annotation files downloaded from the GENCODE portal for mouse genome build GRCm38 release V23. Summary of read and alignment quality were generated using MultiQC v1.12 (Ewels et al., 2016).

#### Differentially expressed genes

Gene expression values were computed as the number of reads aligned per gene, using STAR—quantMode GeneCounts. Raw counts were normalized, and genes with an average read count < 50 across all samples were excluded from the differential analysis. The analysis for differential gene expression was carried out using DESeq2 (Love et al., 2014). A false discovery rate (FDR) cut-off of 0.05 and fold change cut-off of 20% (−0.263 ≤ log2(FC) ≥ +0.263) were used to identify differentially expressed genes.

#### Differential splicing analysis

Alternative splicing events were quantified and classified using rMATS (Shen et al., 2014). Alignment files (BAM) and the GENCODE reference annotation (GTF) for mouse genome build GRCm38 release V23 were used. rMATS classified splicing events into 5 categories: skipped exons, retained introns, mutually exclusive exons, alternative 5’ and 3’ splice sites. An FDR cut-off of 0.05 and an inclusion level difference cut-off of less than −0.2 or greater than 0.2 were used to screen for statistically significant changes.

### Mammalian phenotype ontology analysis

Genes mapping to MPO phenotypes were retrieved from the Mouse Genome Database (Blake et al., 2021). We retrieved a unique set of nodes (n = 15) mapping to the Mammalian Phenotype Ontology terms “impaired adaptive thermogenesis” (MP:0011049) and “abnormal circadian temperature homeostasis” (MP:0011020) and designated these “thermoregulatory nodes” (Smith and Eppig, 2009). We then used the hypergeometric test to evaluate the enrichment of thermoregulatory nodes among nodes with the strongest transcriptional footprints among cold challenge-induced gene sets. The universe was set at the total number of unique nodes included in the HCT intersection analysis (n = 691).

### Consensus transcriptional regulatory network analysis

High confidence transcriptional target (HCT) intersection analysis of gene sets has been previously described (Bissig-Choisat et al., 2021; Chen et al., 2022, p. 1; Ochsner et al., 2020; Zapata et al., 2021). Briefly, consensomes are gene lists ranked according to measures of the strength of their regulatory relationship with upstream signaling pathway nodes derived across numerous independent publicly archived transcriptomic or ChIP-Seq datasets (Ochsner et al., 2019). To generate mouse ChIP-Seq consensomes, we first retrieved processed gene lists from ChIP-Atlas, in which genes are ranked based on their mean MACS2 peak strength across available archived ChIP-Seq datasets in which a given pathway node is the IP antigen (Oki et al., 2018). We then mapped the IP antigen to its pathway node category, class, and family, and organized the ranked lists into percentiles to generate the mouse node ChIP-Seq consensomes (Ochsner et al., 2019). Genes in the 95th percentile of a given node consensome were designated high confidence transcriptional targets (HCTs) for that node and used as the input for the HCT intersection analysis using the Bioconductor GeneOverlap analysis package implemented in R. *P*-values were adjusted for multiple testing by using the method of Benjamini & Hochberg to control the false discovery rate as implemented with the p.adjust function in R, to generate *q*-values (Benjamini and Hochberg, 1995). Evidence for a transcriptional regulatory relationship between a node and a gene set was represented by a larger intersection between the gene set and HCTs for a given node than would be expected by chance after FDR correction (*q* < 0.05).

### Mice and dissection

Immediately after weaning at 3 weeks of age, male wildtype C57BL/6J littermates were housed under thermoneutral (TN, 28°C) or mild cold/room temperature (RT, 22°C) conditions, with an additional cohort of RT-housed mice switched to housing under severe cold (SC, 8°C) conditions for two weeks, starting at 8 weeks of age. All mice were housed in groups of 2-3 per cage, with a 12h light/12h dark cycle, and were provided adequate bedding and nesting material and unrestricted access to water and a standard chow diet (PicoLab 5V5R) throughout the study. 10-week-old ad lib-fed mice were euthanized between zeitgeber time ZT3-ZT6, and interscapular BAT and liver were dissected and weighed after cardiac perfusion with 20 mL PBS. Tissues were snap-frozen in liquid nitrogen and stored at −80°C immediately after dissection. All animal studies were approved by the Baylor College of Medicine Institutional Animal Care and Usage Committee.

### Body Composition

At the end of the study, total fat and lean masses were measured for ad lib-fed mice using an EchoMRI Whole Body Composition Analyzer (Echo Medical Systems) located in the Mouse Metabolic Research Unit at the US Department of Agriculture/Agricultural Research Service (USDA/ARS) Children’s Nutrition Research Center (CNRC) at Baylor College of Medicine. For analysis of body composition, nonfasting glucose levels, and body and tissue weights, statistical analysis was performed using GraphPad Prism software (v8.4). One-way analysis of variance (ANOVA) with Tukey posthoc testing was conducted, and data are presented as mean ± the SEM, with *p* < 0.05 considered statistically significant.

### Blood glucose measurement

Immediately before sacrifice, mice were retroorbitally bled, and nonfasting glucose was measured using a One Touch Ultra 2 glucose meter and test strips (LifeScan).

### Reduced representation bisulfite sequencing

Genomic DNA was extracted from 0.020-0.025g BAT samples obtained from four mice each from TN, RT, and SC groups using a PureLink Genomic DNA Mini Kit (ThermoFisher Scientific) according to manufacturer protocol. Sample libraries were generated from 150 ng genomic DNA and bisulfite conversion was performed using the Ovation RRBS Methyl-Seq System 1-16 with TrueMethyl oxBS kit (Tecan) per manufacturer protocol. After cleanup, library concentration was measured using a Qubit Fluorometer, and fragment distribution was checked using a 2100 Bioanalyzer high sensitivity chip.

Sequencing at a depth of 50-65 million reads per sample was carried out on an Illumina Nextseq 550 System using a NextSeq 500/550 High Output Kit v2.5 (150 Cycles) (Illumina), per manufacturer protocol for 75 bp paired-end sequencing. Libraries were run on an Illumina Nextseq 550 instrument using the MetSeq Primer 1 (NuGEN) mixed with the Read 1 primer (Illumina) according to the Ovation RRBS Methyl-Seq System 1-16 protocol for the first read and the Read 2 primer (Illumina) for the second read, along with standard Illumina indexing primers. Multiplexed samples were spiked with 10% PhiX Control v3 Library calibration control for run quality monitoring. After the run, FASTQ files were generated using Basespace software (Illumina).

#### Data processing

Raw read sequencing quality and adapter contamination were assessed using FastQC v0.11.9 (Andrews, 2010). Overall quality was determined to be satisfactory, and raw reads were aligned to the bisulfite genome using Bismark v0.23.1 (Krueger and Andrews, 2011). The Bismark bisulfite genome was prepared using the GENCODE raw FASTA file for mouse genome build GRCm38 release 23 and the bowtie2 v2.3.5.1 aligner (Langmead and Salzberg, 2012). Alignments were saved as binary format (BAM) files and used to extract methylation calls for CpG, CHG, and CHH contexts using bismark_methylation_extractor. A comprehensive methylation coverage file was also created, detailing the methylation percentage at each base.

#### Differentially methylated regions

Bismark coverage files were used to generate an object compatible with methylKit v1.20.0 (Akalin et al., 2012), summarizing base-level methylation counts for each sample. Only bases with a minimum read coverage of 10 were retained, and then normalized using a scaling factor derived from differences between median coverage distributions between samples. Methylation counts were summarized over the promoter and gene body regions, and differentially methylated promoters and genes were identified using an adjusted *p*-value (*q*-value) cutoff of 0.05.

#### Functional analysis

Functional enrichment analyses were performed on gene lists of interest using WebGestalt (WEB-based GEne SeT AnaLysis Toolkit) with an FDR cutoff of 0.05 (Liao et al., 2019). Minimum number of genes per category was set to 5. Extracted enrichments were then visualized using GoPlot (Walter et al., 2015).

### Histone isolation

One brown adipose tissue (BAT) and one liver datapoint was achieved for each organism from ∼0.1g BAT and ∼0.2g liver. BAT samples were homogenized with 1 mL nuclear isolation buffer (NIB) + 0.3% nonyl phenoxypolyethoxylethanol (NP-40) detergent with appropriate epigenetic inhibitors (sodium butyrate, microcystin-LR, 4-benzenesulfonyl fluoride hydrochloride (AEBSF), and dithiothreitol (DTT)) using a Dounce homogenizer. Nuclei isolation was performed as previously described with optimizations of two NIB + NP-40 incubation steps and two NIB-only washes with a centrifuge duration of 10 minutes at 20,000 *x g* (Holt et al., 2021; Taylor and Young, 2023). Liver samples were homogenized with 1 mL NIB + 1% NP-40 with appropriate epigenetic inhibitors listed above using a Dounce homogenizer. Nuclei isolation was performed as previously described with three NIB-only washes and the centrifuge duration to 10 minutes at 20,000 *x g* (Holt et al., 2021). Acid extraction was performed as previously described (Holt et al., 2021). Isolated histones were resuspended in 85 µL 5% acetonitrile, 0.2% trifluoroacetic acid then histone families were separated by offline high-performance liquid chromatography (HPLC) as described in Holt *et al*. (Holt et al., 2021).

### Histone H3 and H4 mass spectrometry method

#### Histone family offline chromatographic separation

Histone H3 elutes between 42-52 minutes and histone H4 elutes between 34-36.5 minutes with the offline HPLC method used, as previously described (Holt et al., 2021; Taylor and Young, 2023). H3 variants (H3.1, H3.2, H3.3) and H4 fractions were concentrated to dryness with a vacuum centrifuge concentrator (Savant™ SPD131 SpeedVac, Thermo Scientific, Waltham, MA). For histone H3, a standard curve *µg H3 = (Peak area – 2.6558)/14.221* was used to calculate the mass. Each H3 replicate was resuspended with 10 µL ammonium acetate for digestion with 1:10 mass:mass GluC (Sigma-Aldrich #10791156001) at 37 °C for one hour. The digestion was quenched by SpeedVac concentration of the sample to dryness.

#### Preparation of samples for MS and online chromatography

For final dilution, H3 was resuspended to 2 µg/µL using MS buffer A (2% acetonitrile, 0.1% formic acid). For histone H4, dried fractions were diluted using µg H4 = (Peak area – 6.0114)/31.215 to calculate the dilution to 200 ng H4/ µL MS buffer A (2% acetonitrile, 0.1% formic acid). 1 µL diluted histone H3 or H4 was loaded onto a 10 cm, 100 µm inner diameter C3 column (ZORBAX 300SB-C3 300 Å 5 µm) self-packed into fused silica pulled to form a nanoelectrospray emitter. Online HPLC was performed on a Thermo U3000 RSLCnano Pro-flow system. The 70-minute linear gradient using buffer A: 2% acetonitrile, 0.1% formic acid, and B: 98% acetonitrile, 0.1% formic acid is described in **Table 1**. The column eluant was introduced into a Thermo Scientific Orbitrap Fusion Lumos by nanoelectrospray ionization. Static spray voltages of 2400 V for histone H3 or 1800 V for histone H4 and an ion transfer tube temperature of 320 °C were set for the source.

#### Mass spectrometry analysis of histone proteoforms

The orbitrap MS1 experiment used a 60k resolution setting in positive mode. An AGC target of 5.0e5 with 200 ms maximum injection time, three microscans, and scan ranges of 585-640 *m/z* for histone H3 or 700-1400 *m/z* for histone H4 were used. For histone H3, the target precursor selected for MS2 fragmentation included all H3 peaks from GluC peptides and chose the top 6 most abundant *m/z*. ETD fragmentation at 18 ms reaction time, 1.0e6 reagent target with 200 ms injection time was used. MS2 acquisition was performed using the orbitrap with the 30k resolution setting, an AGC target of 5.0e5, a max injection time of 200 ms, a ‘normal’ scan range, and two microscans. For histone H4, an intensity threshold of 1e5 was set for the selection of ions for fragmentation. The target precursor selected for MS2 fragmentation included all major H4 peaks and chose the top 20 most abundant *m/z*. ETD fragmentation at a 14 ms reaction time, 5.0e5 reagent target with 200 ms injection time was used. MS2 acquisition was performed using the orbitrap with the 60k resolution setting, an AGC target of 5.0e5, a max injection time of 200 ms, a ‘normal’ scan range, and three microscans.

### Mass spectrometry data analysis

For histone quantitation, 3-5 animals were analyzed in each of three different housing conditions, giving n=11-15 total for each organ/tissue. Two technical replicates were analyzed per sample and averaged to give the measured value for a minimum of three biological data points per group. Raw files were converted to .mzXML. Data processing was performed by a custom analysis suite as previously described (DiMaggio et al., 2009; Holt et al., 2019, p. 4). TDMS version 7.04 and Interp 2v16 were used for analysis. Search parameters include a 3.4 Da window for MS1, 10.0 ppm tolerance for c and z fragment ions. PTMs considered for histone H4 include fixed N-terminal acetylation and variable K5ac, K8ac, K12ac, K16ac, K20me1/2/3, and K31ac. PTMs considered for histone H3.2 include variable K4me1/2/3, K9ac, K9me1/2/3, K14ac, K18ac, K23ac, K27ac, K27me1/2/3, and K36me1/2/3. This data analysis approach has been used extensively in other studies in our lab to reveal quantitative changes in histone proteoforms (Holt et al., 2019; Jiang et al., 2018, 2019; Taylor and Young, 2023; Wang et al., 2018a, 2018b, 2022). This approach has also been validated against reverse phase protein array based quantitation (Wang et al., 2022). For analyses of BAT versus liver histone modifications, we conducted multiple unpaired t-tests using the false discovery rate approach, and discoveries were determined using the two-stage linear step-up procedure of Benjamini, Krieger and Yekutieli. For all other analyses, Welch’s 2-tailed t-tests or one-way ANOVA with Tukey posthoc testing were used when comparing two or three groups, respectively. *Q-*, *p-*, or *p_adj_-* values less than 0.05 were considered statistically significant.

#### Fold- and absolute change

A fold change cutoff of 1.5 was also used; however, absolute change was also used to inform decisions on biological significance. Unlike many proteomic methods, we accurately measure absolute change. In our experience, it is not uncommon for large absolute changes to result in small-fold changes yet have profound biological significance. For example, a histone PTM that decorates 50% of the entire genome that increases to 70% results in a change to an astounding 20% of the genome but is less than 1.5-fold. Thus, consideration of both fold change and absolute change is insightful and represented as independent metrics.

### Experimental Design and Statistical Rationale

Unless otherwise stated, experiments were designed with an N of 3-5 per group (BAT or liver from mice housed at TN, RT, and SC). The data obtained was interpreted collectively for a robust understanding of the system. Rigor was enhanced through multiple complimentary methods to understand epigenetic changes during BAT thermogenesis. Various parametric statistics and corrections were used and are described above. Generally, one-way ANOVA with Tukey post-hoc testing or Welch’s two-tailed t-tests were used with *p* < 0.05 considered statistically significant. *P_adj_*or *q* values are calculated where appropriate, namely with consensus transcriptional regulatory network and RRBS analyses and comparisons between tissue histone modifications at each temperature.

## Acknowledgments

B.C.T. and N.L.Y. were supported by NIH Grants R01 GM139295, P01 AG066606, RF1 AG074540, R01 CA193235, R01 CA276663, and R56 HG012206. L.H.S. and A.M.N.- A. were supported by USDA-ARS Cooperative Agreement 3092-51000-064-005-S, and A.M.N.A. was additionally supported by American Heart Association Career Development Award 18CDA34110137 and a Texas Children’s Hospital Pediatrics Pilot Award. M.D., H.K.Y., and G.E.Z. and N.R.M. were supported by United States Department of Agriculture (USDA/ARS) Cooperative Agreement No. 58-3092-0-001. H.K.Y. was additionally supported by a Duncan NRI Zoghbi Scholar Award to H.K.Y. S.A.O. and N.J.M. were supported by NIDDK Information Network Grant U24DK097771. The contents of this work are solely the responsibility of the authors and do not necessarily represent the official views of the USDA.

## Author Contributions

B.C.T., H.K.Y., N.J.M., N.L.Y., and A.M.N.-A. conceptualization; B.C.T., L.H.S., M.D., H.K.Y., S.A.O., G.E.Z., N.R.M., N.J.M., N.L.Y., and A.M.N.-A. methodology and investigation; B.C.T., M.D., H.K.Y., S.A.O., N.J.M., and A.M.N.-A. formal analyses; B.C.T., M.D., N.R.M., N.J.M., N.L.Y. and A.M.N.-A. writing-original manuscript; N.L.Y. and A.M.N.-A. supervision and funding acquisition.

## Data Availability

RRBS data are deposited in GEO (accession number GSE234588). Histone H3.2 and H4 mass spectrometry raw data are uploaded to the MassIVE repository (ftp://massive.ucsd.edu/MSV000092105/). All other data described in this manuscript are contained within the manuscript or as associated supplementary material.

## Supplemental data

This article contains supplemental data.

## Abbreviations

ACN: Acetonitrile
AEBSF: 4-benzenesulfonyl fluoride hydrochloride
AGC: Automatic gain control
ANOVA: One-way analysis of variance
BAT: Brown adipose tissue
C3: 3 carbon chain
CBP: CREB-Binding Protein
ChIP-Seq: Chromatin immunoprecipitation sequencing
DEG: Differentially expressed genes
DMP: Differentially methylated promoters
DNA: Deoxyribonucleic acid
DNMT1: DNA methyltransferase 1
DNMT3B: DNA methyltransferase 3B
DNL: Day night average sound level
DTT: Dithiothreitol
ETD: Electron transfer dissociation
EZH2: Enhancer of zeste homolog 2
FA: Formic acid
FC: Fold change
FCOR: FOXO1-corepressor
FDR: False discovery rate
FOXO1: Forkhead Box O1
G9a/GLP (EHMT2/EHMT1): Euchromatic histone-lysine N-methyltransferase 2/ Euchromatic histone-lysine N-methyltransferase 1
GluC: Staphylococcus aureus Protease V8
HCT: High confidence transcriptional target
HDAC3: Histone deacetylase 3
HPLC: High-performance liquid chromatography
K: Lysine
Kat8: Lysine Acetyltransferase 8
Kdm2b: Lysine (K)-specific demethylase
2B: Kdm3a Lysine demethylase 3A
Kdm5b: Lysine-specific demethylase
5B: Kdm5c Lysine demethylase 5C
Kdm8 (JMJD5): Lysine demethylase 8
LC-MS/MS: Liquid chromatography tandem mass spectrometry
MACS2: Model-based analysis of ChIP-seq
MS: Mass spectrometry
MS2: Tandem mass spectrometry or mass spectrum
mzXML: File format, eXtensible markup language
NAD+: Nicotinamide adenine dinucleotide
NIB: Nuclei isolation buffer
NP-40: Nonyl phenoxypolyethoxylethanol
NST: Nonshivering thermogenesis
OR: Odds ratio
P300: E1A-Binding Protein, 300-KD
PRC2: Polycomb repressive complex 2
PTM: Post-translational modification
RNA-Seq: RNA sequencing
RRBS: Reduced representation bisulfite sequencing
RT: Room temperature
SAM: S-adenosyl methionine SIRT1 Sirtuin-1
SC: Severe cold
SEM: Standard error of the mean
Setd7: Histone-lysine N-methyltransferase SETD7
SIRT1: Sirtuin-1
SMYD3: SET and MYN-domain containing 3
SRA: Sequence Read Archive
SUV39H1: Histone-lysine N-methyltransferase SUV39H1
SUV39H2: Histone-lysine N-methyltransferase SUV39H2
TFA: Trifluoroacetic acid
TN: Thermoneutral
UCP1: Uncoupling protein 1
WAT: White adipose tissue

---

**Table S1.**
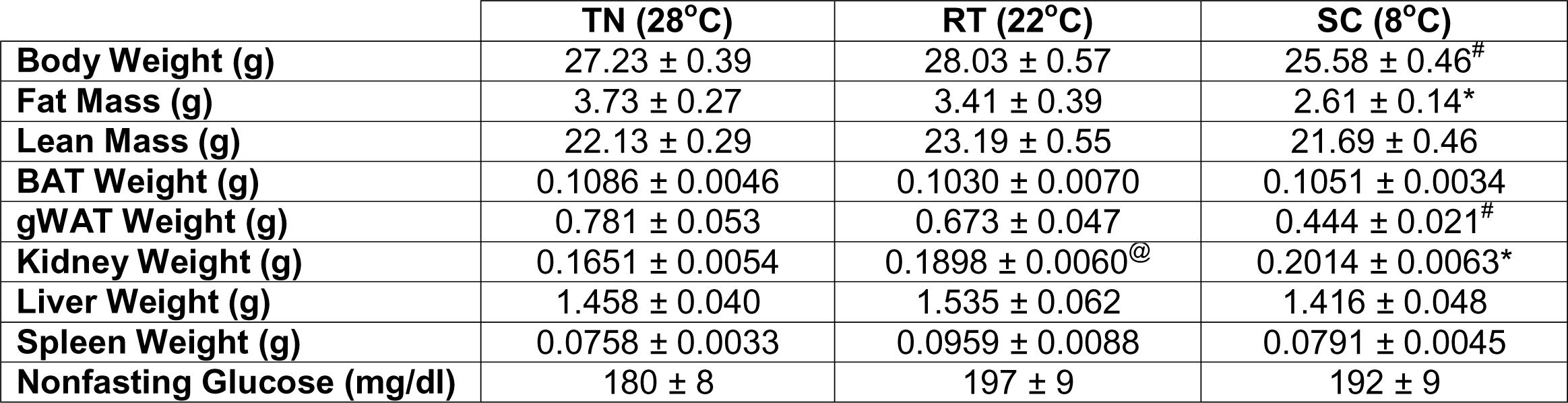
Average body weights, fat mass, lean mass, tissue weights, and nonfasting glucose of male C57BL/6J mice after exposure to thermoneutral or chronic cold housing temperatures. Results shown are mean ± SEM. \**p* <0.05 for SC vs TN, ^#^*p* <0.05 for SC vs RT and TN, and ^@^*p* <0.05 for RT vs TN. TN: thermoneutral, RT: room temperature, SC: severe cold.

|  | **TN (28°C)** | **RT (22°C)** | **SC (8°C)** |
| --- | --- | --- | --- |
| **Body Weight (g)** | 27.23 ± 0.39 | 28.03 ± 0.57 | 25.58 ± 0.46<sup>#</sup> |
| **Fat Mass (g)** | 3.73 ± 0.27 | 3.41 ± 0.39 | 2.61 ± 0.14<sup>*</sup> |
| **Lean Mass (g)** | 22.13 ± 0.29 | 23.19 ± 0.55 | 21.69 ± 0.46 |
| **BAT Weight (g)** | 0.1086 ± 0.0046 | 0.1030 ± 0.0070 | 0.1051 ± 0.0034 |
| **gWAT Weight (g)** | 0.781 ± 0.053 | 0.673 ± 0.047 | 0.444 ± 0.021<sup>#</sup> |
| **Kidney Weight (g)** | 0.1651 ± 0.0054 | 0.1898 ± 0.0060<sup>@</sup> | 0.2014 ± 0.0063<sup>*</sup> |
| **Liver Weight (g)** | 1.458 ± 0.040 | 1.535 ± 0.062 | 1.416 ± 0.048 |
| **Spleen Weight (g)** | 0.0758 ± 0.0033 | 0.0959 ± 0.0088 | 0.0791 ± 0.0045 |
| **Nonfasting Glucose (mg/dl)** | 180 ± 8 | 197 ± 9 | 192 ± 9 |

**Table S2.**
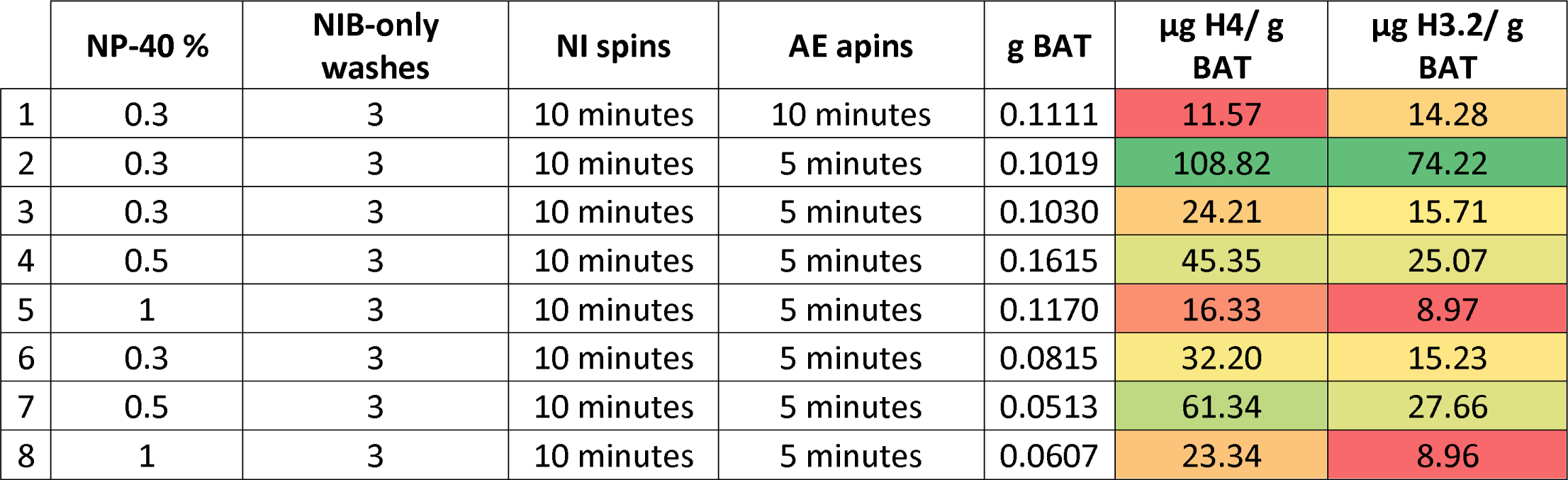
Results of method optimization of histone isolation from BAT. Color scheme indicates the highest yield of histones in green (2) and the lowest yield of histones in red. Mass histone is calculated using offline HPLC peak area and standard curve.

**Table S3.**
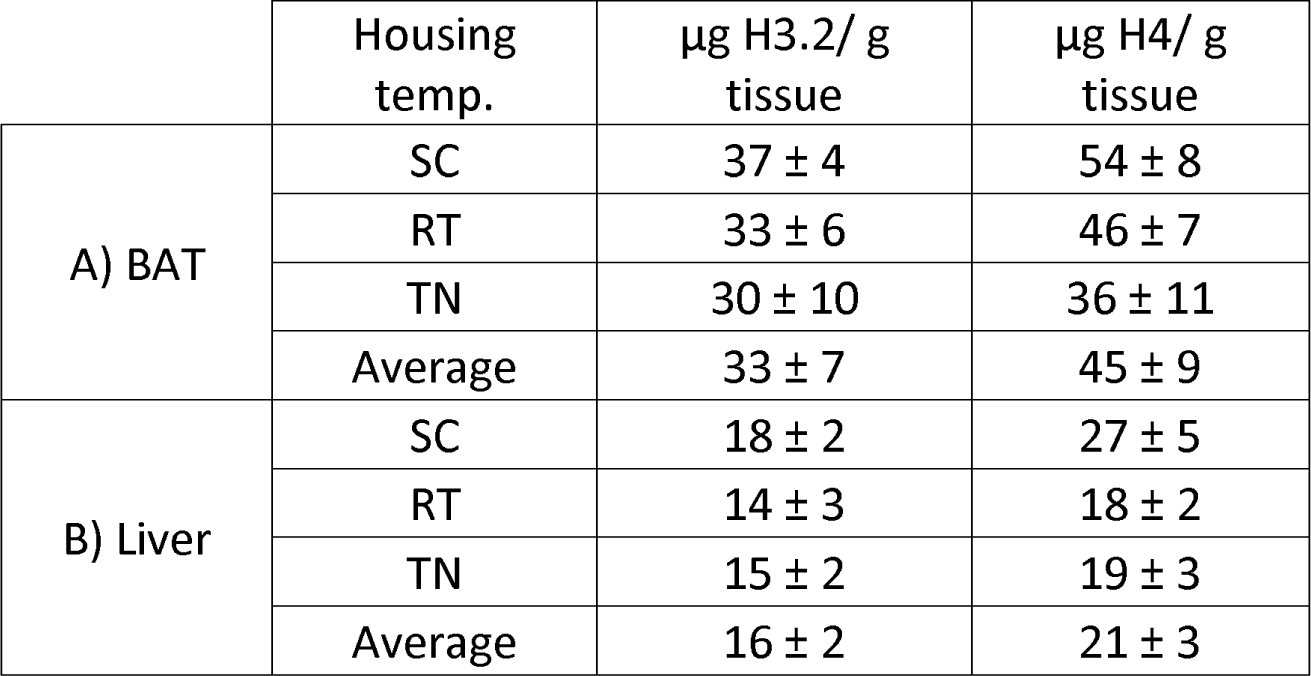
Mass of histone H3.2 and H4 obtained from **(a)** BAT and **(b)** liver at different housing temperatures. Mass histone is calculated using offline HPLC peak area and standard curve. TN: thermoneutral; RT: room temperature; SC: severe cold.

