## Supplemental Table 1 for "Histone proteoform analysis reveals epigenetic changes in adult mouse brown adipose tissue in response to cold stress"

***Table S1.*** Average body weights, fat mass, lean mass, tissue weights, and nonfasting glucose of male C57BL/6J mice after exposure to thermoneutral or chronic cold housing temperatures. Results shown are mean ± SEM. **p* <0.05 for SC vs TN, #*p* <0.05 for SC vs RT, and @*p* <0.05 for RT vs TN. TN: thermoneutral, RT: room temperature, SC: severe cold.

| | **TN (28$^{o}$C)** | **RT (22$^{o}$C)** | **SC (8$^{o}$C)** |
|---|---|---|---|
| **Body Weight (g)** | 27.23 ± 0.39 | 28.03 ± 0.57 | 25.58 ± 0.46*# |
| **Fat Mass (g)** | 3.73 ± 0.27 | 3.41 ± 0.39 | 2.61 ± 0.14* |
| **Lean Mass (g)** | 22.13 ± 0.29 | 23.19 ± 0.55 | 21.69 ± 0.46 |
| **BAT Weight (g)** | 0.1086 ± 0.0046 | 0.1030 ± 0.0070 | 0.1051 ± 0.0034 |
| **gWAT Weight (g)** | 0.781 ± 0.053 | 0.673 ± 0.047 | 0.444 ± 0.021*# |
| **Kidney Weight (g)** | 0.1651 ± 0.0054 | 0.1898 ± 0.0060@ | 0.2014 ± 0.0063* |
| **Liver Weight (g)** | 1.458 ± 0.040 | 1.535 ± 0.062 | 1.416 ± 0.048 |
| **Spleen Weight (g)** | 0.0758 ± 0.0033 | 0.0959 ± 0.0088 | 0.0791 ± 0.0045 |
| **Nonfasting Glucose (mg/dl)** | 180 ± 8 | 197 ± 9 | 192 ± 9 |
