## Supplemental Table 2 for "Histone proteoform analysis reveals epigenetic changes in adult mouse brown adipose tissue in response to cold stress"

***Table S2.*** *Results of method optimization of histone isolation from BAT.* Color scheme indicates the highest yield of histones in green (2) and the lowest yield of histones in red. Mass histone is calculated using offline HPLC peak area and standard curve.

| | NP-40 % | NIB-only washes | NI spins | AE apins | g BAT | µg H4/ g BAT | µg H3.2/ g BAT |
|---|---|---|---|---|---|---|---|
| 1 | 0.3 | 3 | 10 minutes | 10 minutes | 0.1111 | 11.57 | 14.28 |
| 2 | 0.3 | 3 | 10 minutes | 5 minutes | 0.1019 | 108.82 | 74.22 |
| 3 | 0.3 | 3 | 10 minutes | 5 minutes | 0.1030 | 24.21 | 15.71 |
| 4 | 0.5 | 3 | 10 minutes | 5 minutes | 0.1615 | 45.35 | 25.07 |
| 5 | 1 | 3 | 10 minutes | 5 minutes | 0.1170 | 16.33 | 8.97 |
| 6 | 0.3 | 3 | 10 minutes | 5 minutes | 0.0815 | 32.20 | 15.23 |
| 7 | 0.5 | 3 | 10 minutes | 5 minutes | 0.0513 | 61.34 | 27.66 |
| 8 | 1 | 3 | 10 minutes | 5 minutes | 0.0607 | 23.34 | 8.96 |
