## Supplemental Table 3 for "Histone proteoform analysis reveals epigenetic changes in adult mouse brown adipose tissue in response to cold stress"

***Table S3.*** *Mass of histone H3.2 and H4 obtained from* ***(a)*** *BAT and* ***(b)*** *liver at different housing temperatures.* Mass histone is calculated using offline HPLC peak area and standard curve. TN: thermoneutral; RT: room temperature; SC: severe cold.

| | Housing temp. | µg H3.2/ g tissue | µg H4/ g tissue |
|---|---|---|---|
| A) BAT | SC | 37 ± 4 | 54 ± 8 |
| | RT | 33 ± 6 | 46 ± 7 |
| | TN | 30 ± 10 | 36 ± 11 |
| | Average | 33 ± 7 | 45 ± 9 |
| B) Liver | SC | 18 ± 2 | 27 ± 5 |
| | RT | 14 ± 3 | 18 ± 2 |
| | TN | 15 ± 2 | 19 ± 3 |
| | Average | 16 ± 2 | 21 ± 3 |
